# The Protein Design Archive (PDA): insights from 40 years of protein design

**DOI:** 10.1101/2024.09.05.611465

**Authors:** Marta Chronowska, Michael J. Stam, Derek N. Woolfson, Luigi F. Di Constanzo, Christopher W. Wood

## Abstract

The field of protein design has changed dramatically over the last 40 years, with a range of methods developing from rational design to more recent data-driven approaches. While considerable insight could be gained from analysing designed proteins, there is no single resource that brings together all the relevant data, making it difficult to identify current challenges and opportunities within the field.

Here we present the Protein Design Archive, a website and database of designed proteins. Using the database, we performed systematic analysis that reveals a rapid increase in the number and complexity of designs over time and uncovers biases in their amino-acid usage and secondary structure content. Finally, sequence- and structure-based analysis demonstrates the breadth and novelty of designs within the archive. The PDA will be a valuable resource for guiding development of protein design, helping the field reach its potential.

The PDA is available freely and without registration at https://pragmaticproteindesign.bio.ed.ac.uk/pda/.

## Introduction

Natural proteins are an incredibly diverse and versatile group of biomacromolecules with a wide range of functions and potential applications as materials ^1,2^, biosensors ^3^, and therapeutics ^4^. In addition, as enzymes, their remarkable product stereospecificity, efficiency, and catalytic activity at ambient temperature makes them valuable for the production of pharmaceutical, specialty, and bulk chemicals ^5^. Nonetheless, nature has explored only a minuscule subset of the incredibly vast sequence and structural space available to protein chains ^6,7^. The field of protein design attempts to delve into the unexplored regions of protein space, to deepen our understanding of the protein sequence-to-structure/function relationships, and to develop novel proteins to tackle industrial and societal challenges.

A range of approaches have been applied to design proteins with the earliest attempts in the 1980s (table 1). This “infancy” period (∼1980 – 1990) was characterised by the *minimal* approach to protein design, relying on fundamental physicochemical properties of proteins, such as the assembly of amphipathic α helices into bundles ^8^. In the “adolescent” period (∼1990 – 2010) of protein design, *minimal* approaches were extended to *rational*, or *fragment-based and bioinformatically informed* strategies ^7^. In these, the basic patterns of polar and hydrophobic residues used in minimal design were enhanced with sequence-to-structure relationships gained from evolutionary, biochemical, bioinformatics, or empirical studies ^9–11^.

**Table 1.**
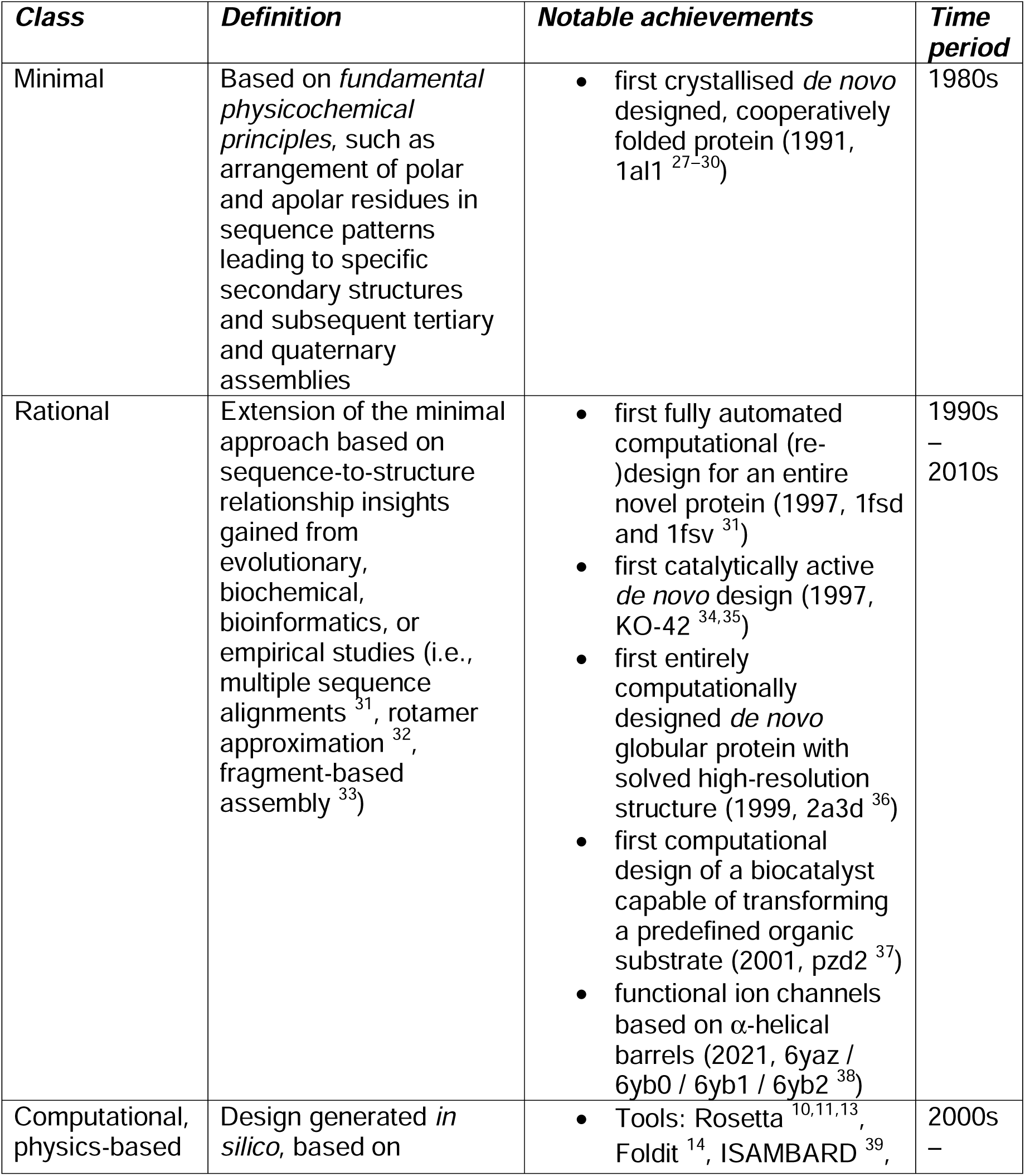
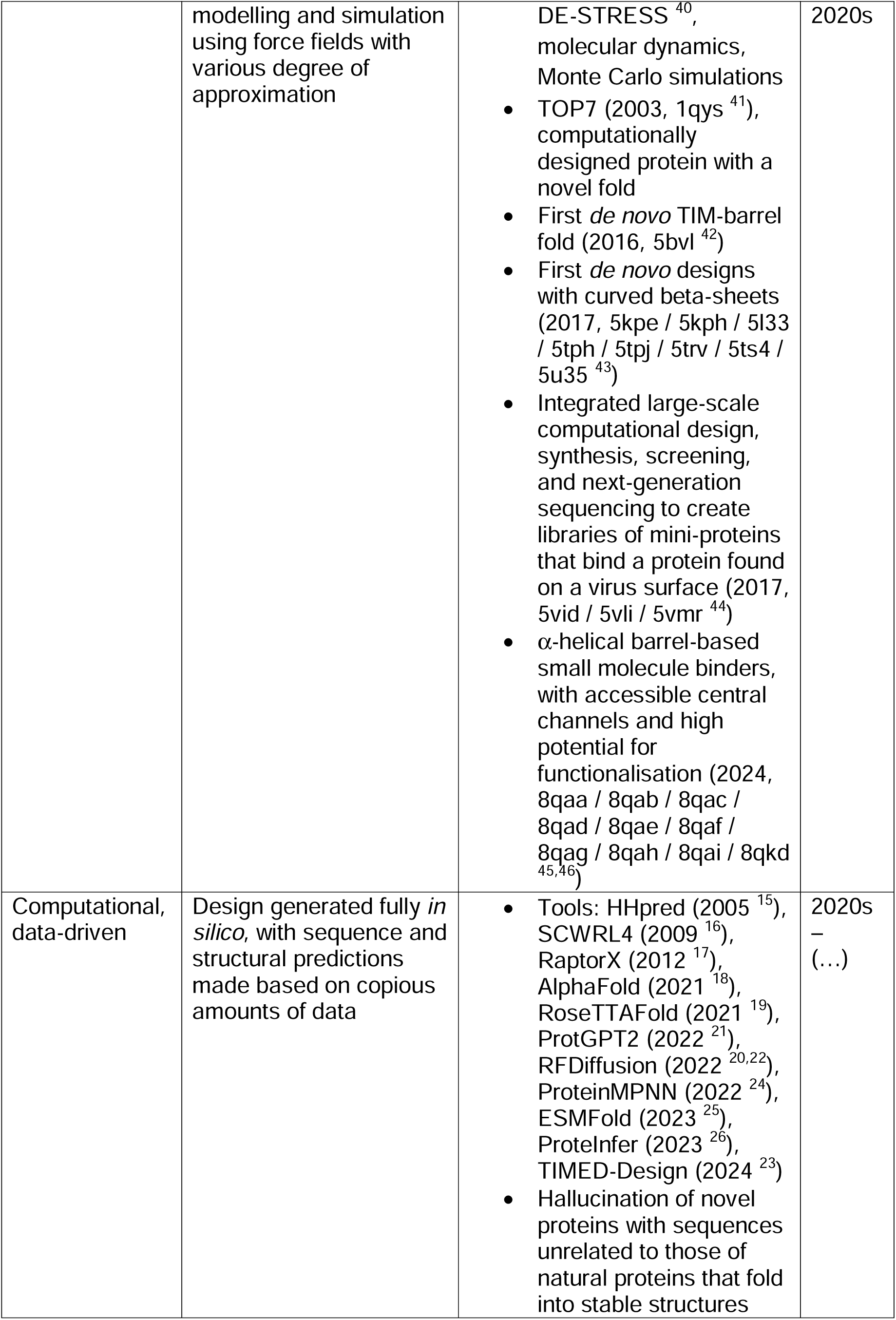
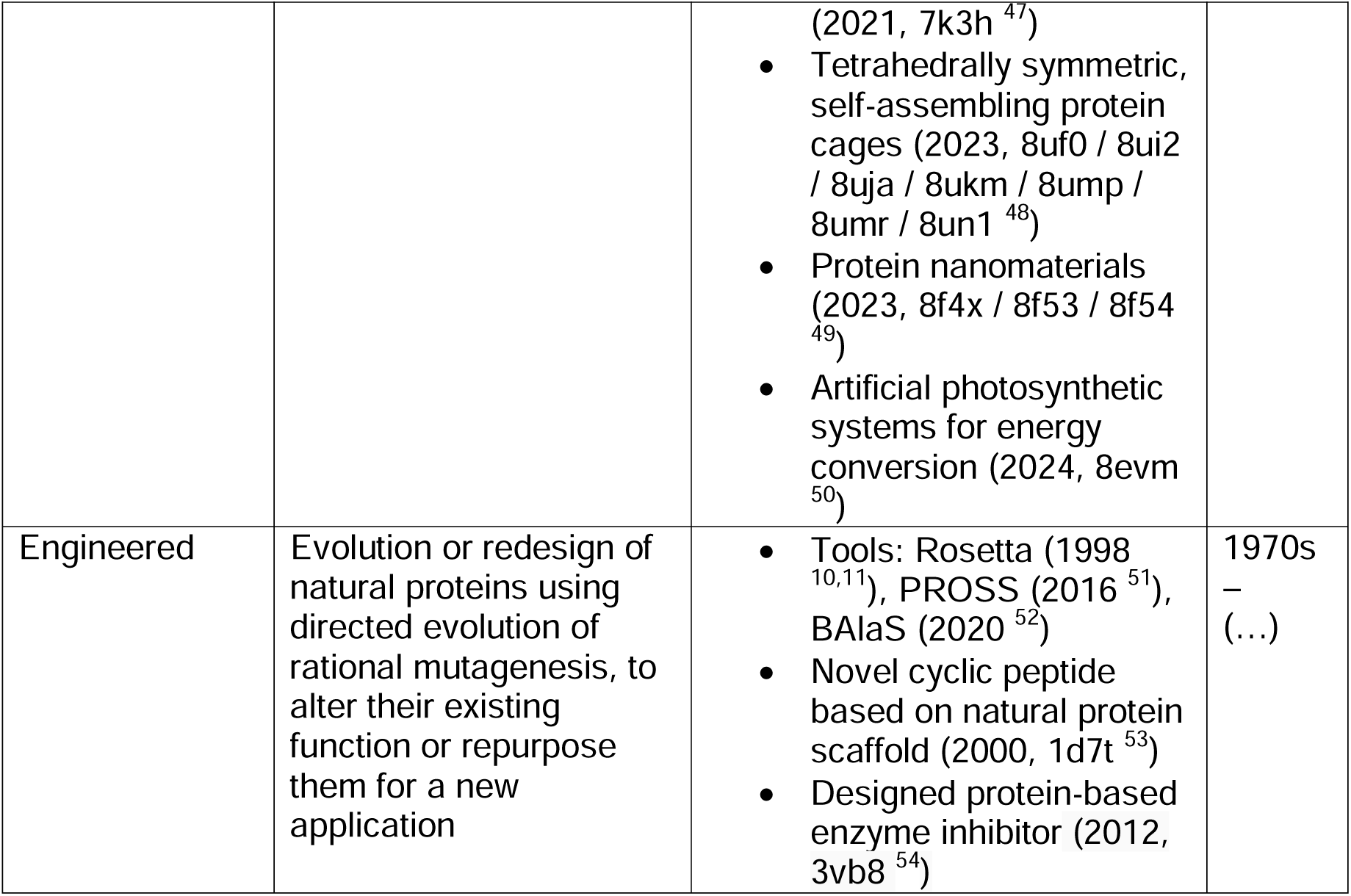
Classification of protein designs based on their development strategy, including definition, notable achievements, and period of popularity.

It has been said that the field of protein design has “come of age” over the last decade (∼2010 – present) ^12^, largely due to rapid advancements in computational approaches. These approaches can be divided into two main categories: *physics-based* methods and *data-driven* methods. While all methods generate proteins primarily *in silico*, physics-based approaches simulate atomic-level interactions using force fields with varying degrees of approximation ^10,11,13,14^. In contrast, *data-driven* methods predict sequence and structure probabilities by identifying subtle patterns from vast amounts of data ^15–17^, with deep learning-based methods gaining popularity over recent years ^18–26^.

Over time, many attempts have been made to capture progress in the ever-evolving field of protein design. While many excellent reviews exist ^7,9,12,55–59^, rapid progress ensures that no single review can comprehensively cover the breadth of the field, and these efforts will become increasingly difficult going forward due to the accelerated rise in the number of designed proteins.

We have developed the Protein Design Archive (PDA), a web application and associated database that enables users to explore designed proteins that have been structurally characterised. For this first version, the designs in the archive are hand-curated to ensure that the user is presented with high-quality information and has as much context about the design as possible, while laying the foundations required for automatic curation and classification of entries in the future. A simple user interface enables browsing as well as filter-based searching, and export of data for further use. Analysis of the designs within the database reveals present biases in their sequence and structure. Finally, sequence and structural analysis provides insights into the novelty of designed proteins.

## Results

### Database

At the time of publishing this paper, the PDA database contains 1,450 protein design structures (fig. 1 and 2). These consist of proteins of synthetic origin deposited in the RCSB PDB database, a wwPDB partner and archive for macromolecular information, supplemented manually based on knowledge of the field and literature research. For each design we collected publication, structural, and sequence information.

**Figure 1:**
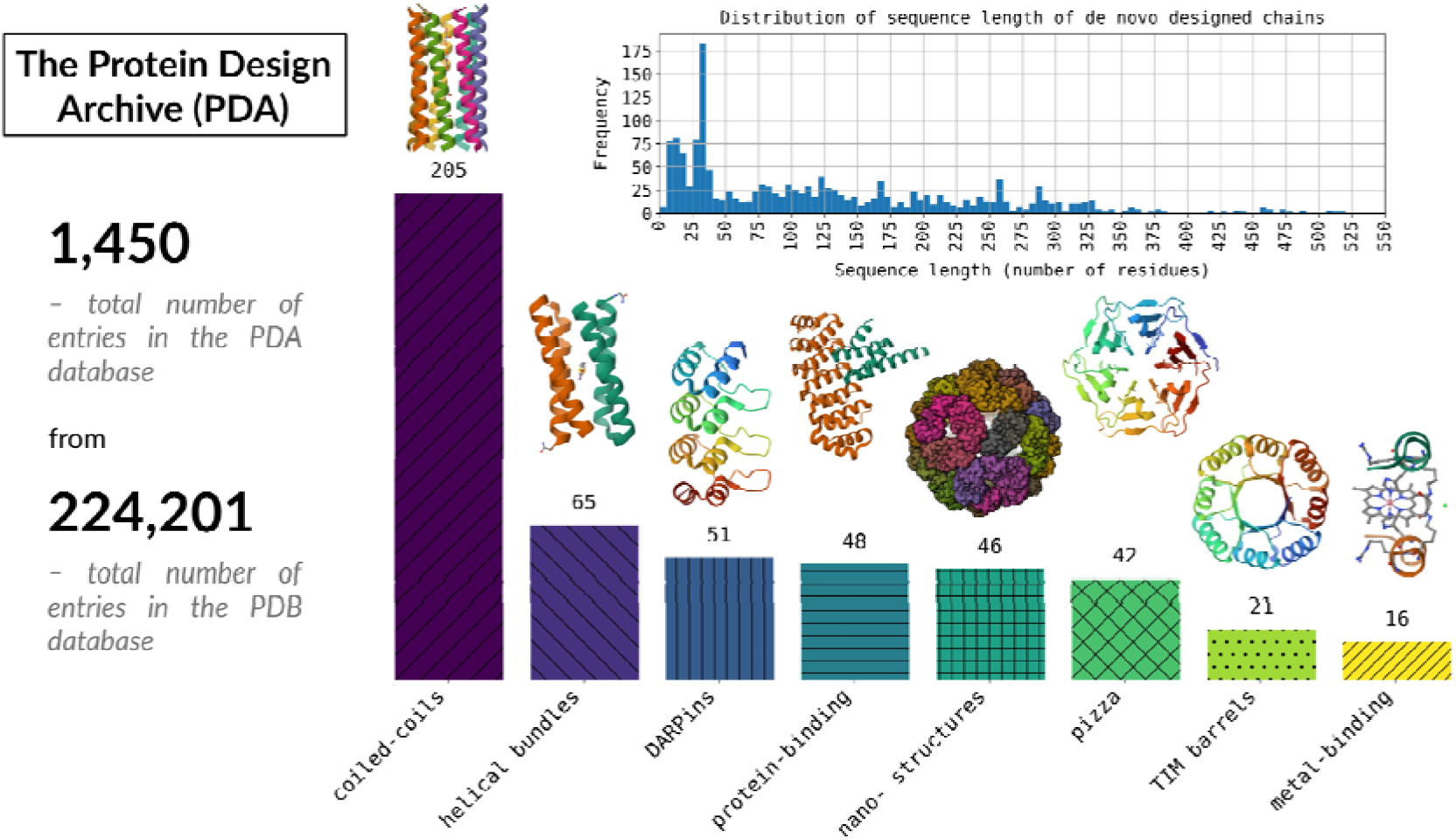
Overview of data in the PDA database distinguishing selected common types of protein designs – coiled coils, helical bundles, DARPins, protein binders, nano(-tube, -wire) structures, β-propellor proteins, TIM barrels, and metal binders – with their number of entries in the database, and example structures (from left to right: 4pna, 1jm0, 2jab, 4jw2, , 8f54, 6rli, 6wvs, 1pyz).

**Figure 2:**
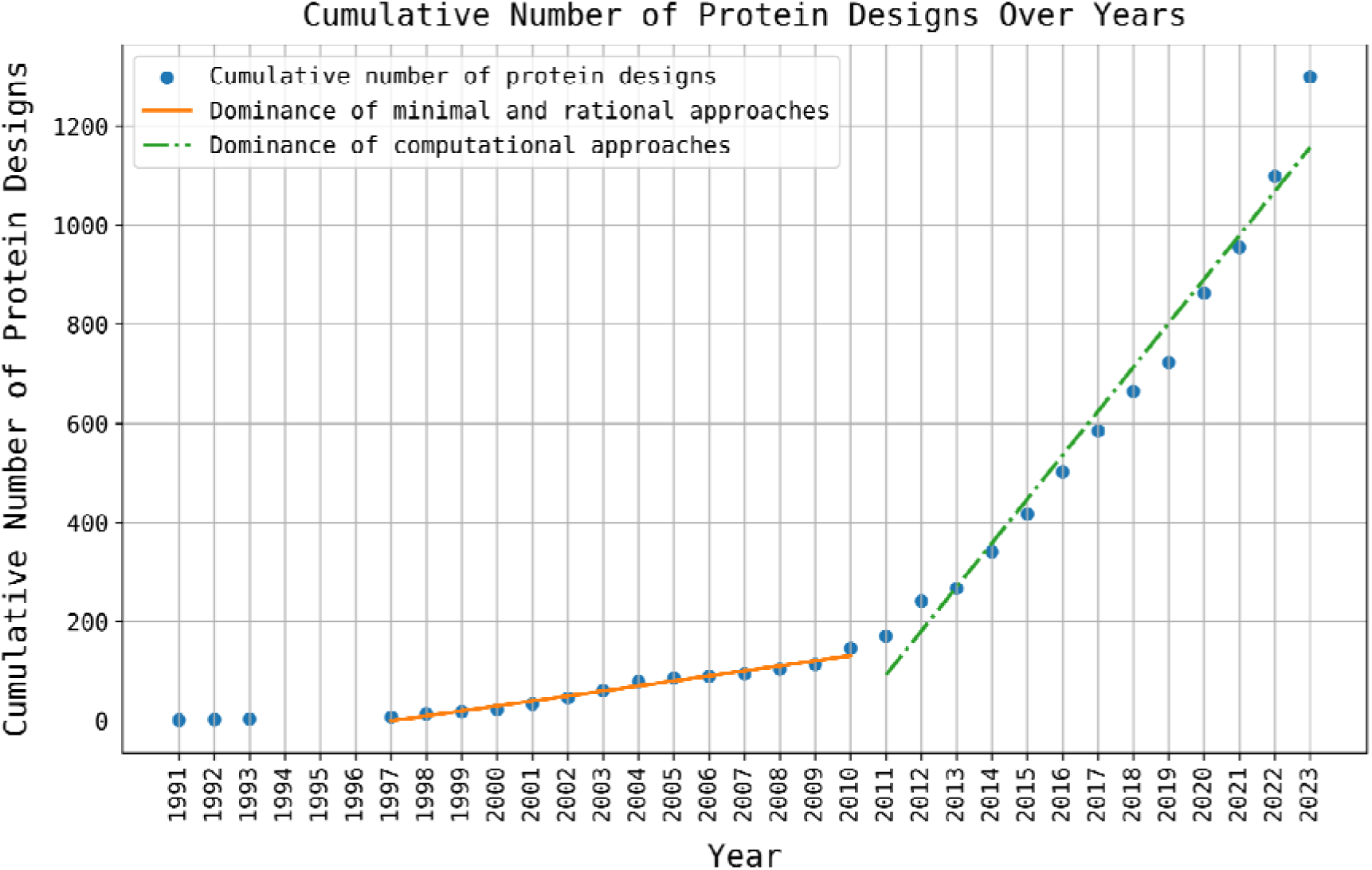
Growth of designed protein structures. Two regions have been distinguished based on the dominant approaches to protein design over these time periods: minimal and rational (1997 – 2010), and computational (from 2010).

Figure 2 shows the growth of protein designs in the PDA over time. Two distinct regions can be identified in this plot distinguished by the dominant approach to protein design at different times: minimal and rational (from the first routine protein design experiments in 1997 to 2010), and computational (from 2010 until now). Linear fits of both time periods (table S1), reveal a 9-fold increase in the rate of release of new protein designs, from an average of 10 to 89 new designed structures per year. The doubling of designs deposited every year from 2020 compared to previous years (table S2) might be an early indication that potentially we have entered another distinct period of deep learning-based protein design, which may bring an unprecedented number of designs and resulting structures, although this remains to be determined.

Among experimental methods for structural determination of the current entries contained in PDA, X-ray crystallography (1,150 structures) and NMR spectroscopy (235 structures) remain the primary methods for designed protein structure determination, while cryo-electron microscopy (cryo-EM; 64 structures) is gaining popularity rapidly. Of the 1,450 entries, 41% are oligomeric assemblies with either point group symmetry or helical symmetry.

### User Interface

The interface of the PDA has been structured to enable free browsing for learning and inspiration as well as targeted searches by the user. The landing page of the PDA database, at https://pragmaticproteindesign.bio.ed.ac.uk/pda/, serves as a summary page. It is arranged as a timeline plot showing all designs matching the current filtering criteria (fig. 3A). Hovering over a point on the plot highlights it and displays the entry’s PDB code, and the point can be clicked to navigate to this design’s details page.

**Fig. 3.**
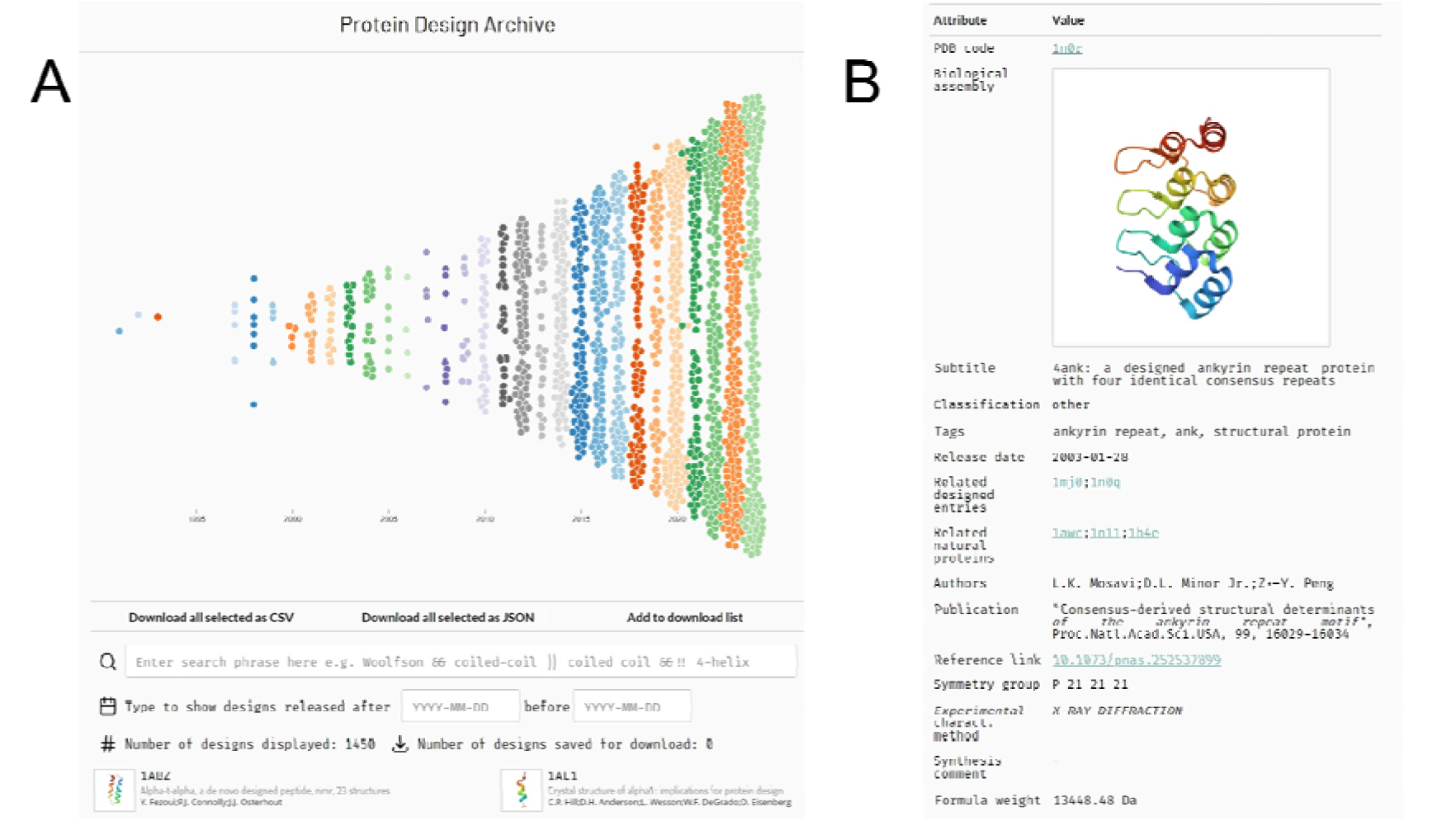
User interface of the PDA website. A) Home page view. The designs are arranged as points on a timeline plot from year 1991 (release of 1al1) to 2024 (time of publication of this paper), showing the rapid growth of designs across the years. B) Design details page. Main information is summarised in an index card-style.

The currently allowed filtering criteria are based on text, release date ranges, and sequence and structure similarity to natural proteins. The number of designs corresponding to the current filtering criteria is displayed below the search section, simultaneously informing of the total number of designs in the database, if no criteria are imposed. Users can export all or a selected subset of designs in a CSV or JSON file format.

Each design has a dedicated details page (fig. 3B), with information about the entry, a static 2D image of the biological assembly, a subtitle, tags, the release date, publication citation and link, authors, protein structures related by sequence or structure, formula weight, a synthesis comment, an interactive structure viewer (using NGL viewer ^60,61^) and a short “Description” section, based on the publication’s abstract.

### Analysis of designed proteins against the PDB

Properties of designed proteins in the PDA database were analysed and compared to protein structures from the PDB. Figure 4A and 4B show heatmaps of the amino acid and secondary structure proportions of designed proteins over time, along with a comparison against the PDB. Both heatmaps are coloured by the percentage increase (red) or decrease (blue) in the proportions compared to those of the PDB. One key observation from figure 4A is that the proportions of glutamate (E), lysine (K) and non-standard residues (X), are over twice as high in the period 1995 – 2000 than in the PDB. Additionally, other amino acids are underrepresented in this period, such as valine (V) at 1.2% compared to 7.0% in the PDB, and cysteine (C) at 0.8% compared to 1.5% in the PDB. Across the different time periods, some of the amino-acid proportions in designed proteins have become closer to those found in the PDB, for example, designs had 6.6% of valine (V) in 2015 – 2020 and 6.7% in 2020 – 2025. However, there are still significant differences between the amino-acid composition of designed proteins and the PDB in the most-recent time period (2020 – 2025), with both alanine (A) and glutamate (E) at nearly twice the level of the PDB, and other amino acids such as phenylalanine (F), cysteine (C) and threonine (T) are underrepresented.

**Figure 4:**
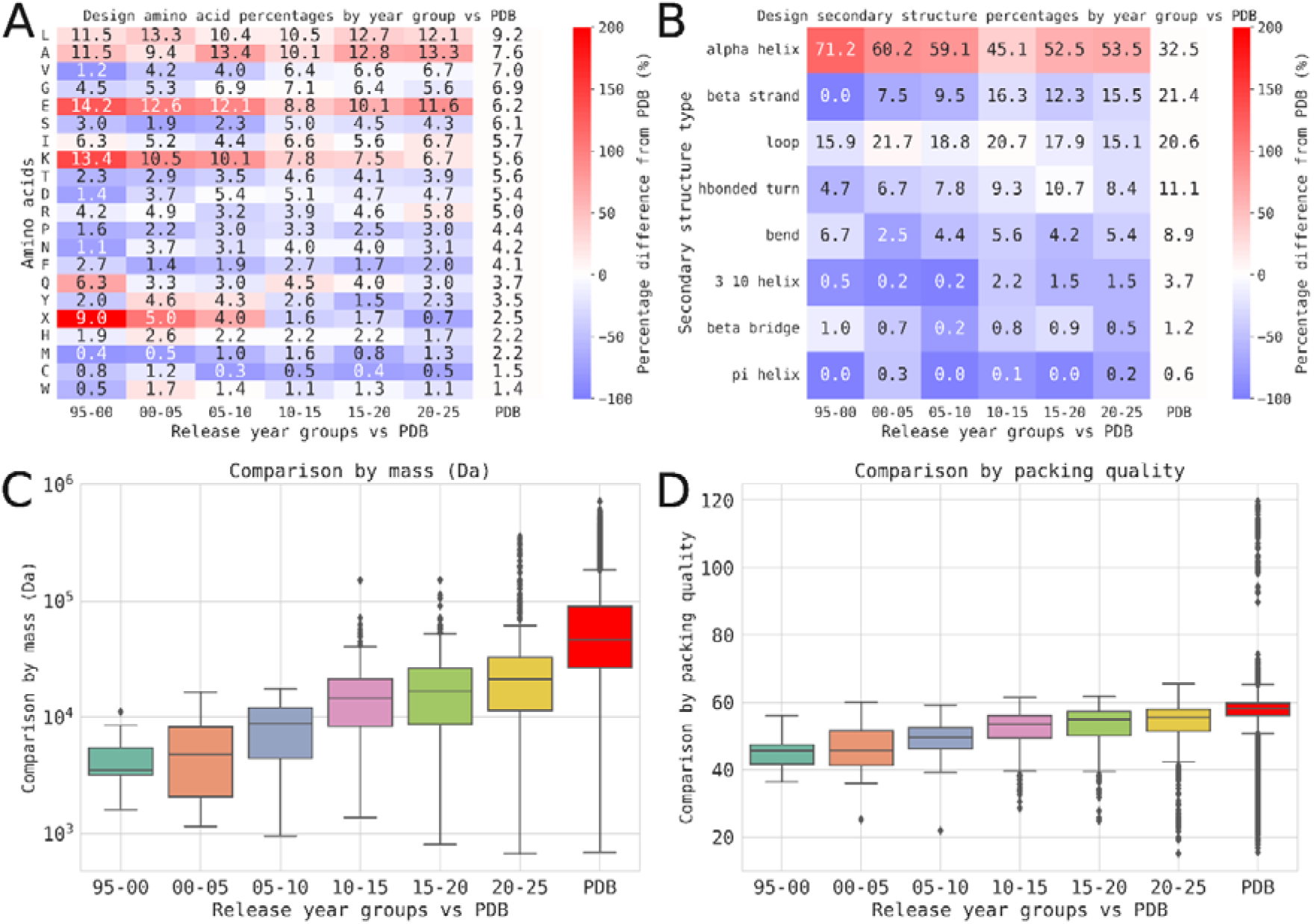
A) A heat map showing amino acid proportions for designs vs the PDB by release year group. The heat map is coloured by the percentage increase (red) or decrease (blue) against the PDB. B) A heat map showing secondary structure proportions for designs vs the PDB by release year group. The heat map is coloured by the percentage increase (red) or decrease (blue) against the PDB. C) Box plots of mass in Daltons (Da) for the designed proteins vs the PDB by release year group. The y-axis is on a logarithmic scale. D) Box plots of packing density of the designed proteins vs the PDB by release year group.

In addition to this, figure 4B shows that α-helical secondary structure is overrepresented in designed proteins across all time periods compared to the PDB, while other secondary structure types such as β strand, β bridge and 3_10_ helices are underrepresented. For example, in the time periods 2015 – 2020 and 2020 – 2025 the proportion of α helices in designs is 52.5% and 53.5%, respectively, whereas in the PDB it is 32.5%, while in 2015 – 2020 the proportion of β strands is 12.3%, compared to 15.5% in 2020 – 2025 and 21.4% in the PDB. This analysis demonstrates that designed proteins are biased towards using certain residues and secondary structure types in comparison to the PDB as a whole.

Following the analysis of composition, the DE-STRESS software ^40^ was used to generate a range of other properties for designed proteins and the wider PDB. Figure 4C displays box plots of mass, in Daltons (Da), for the designed proteins across time and for the PDB. This plot shows a clear trend over time, with designed proteins becoming increasingly larger. However, it also shows that on average, proteins are not being designed as large as those in the PDB more widely. Additionally, the packing quality of designed proteins and the PDB were calculated in figure 4D by using the mean packing density, obtained using DE-STRESS, where the packing density of a non-hydrogen atom is defined as the number of non-hydrogen atoms within a 7 Å radius ^39^. From these box plots, we observe a similar trend over time, where the packing quality of designed proteins has been steadily increasing, although the packing quality of these proteins is still currently lower on average than most proteins in the PDB.

### MMseqs2 and Foldseek analysis of sequence and structure similarity

To gain insight into the novelty of the designed proteins, a sequence similarity search using MMseqs2 ^62^ and a structure similarity search using Foldseek ^63^ was performed. We analysed how designed chains compare to natural proteins (DvP-type search,) and how they are related to one another (DvD-type search).

For each metric and every design, we find the highest scoring query–target pair. Full list of analysed metrics is reported in the supplementary material (fig. S1 and S2), and summary of the results for sequence-based bit scores and structure-based LDDT is shown in figure 5.

**Fig. 5:**
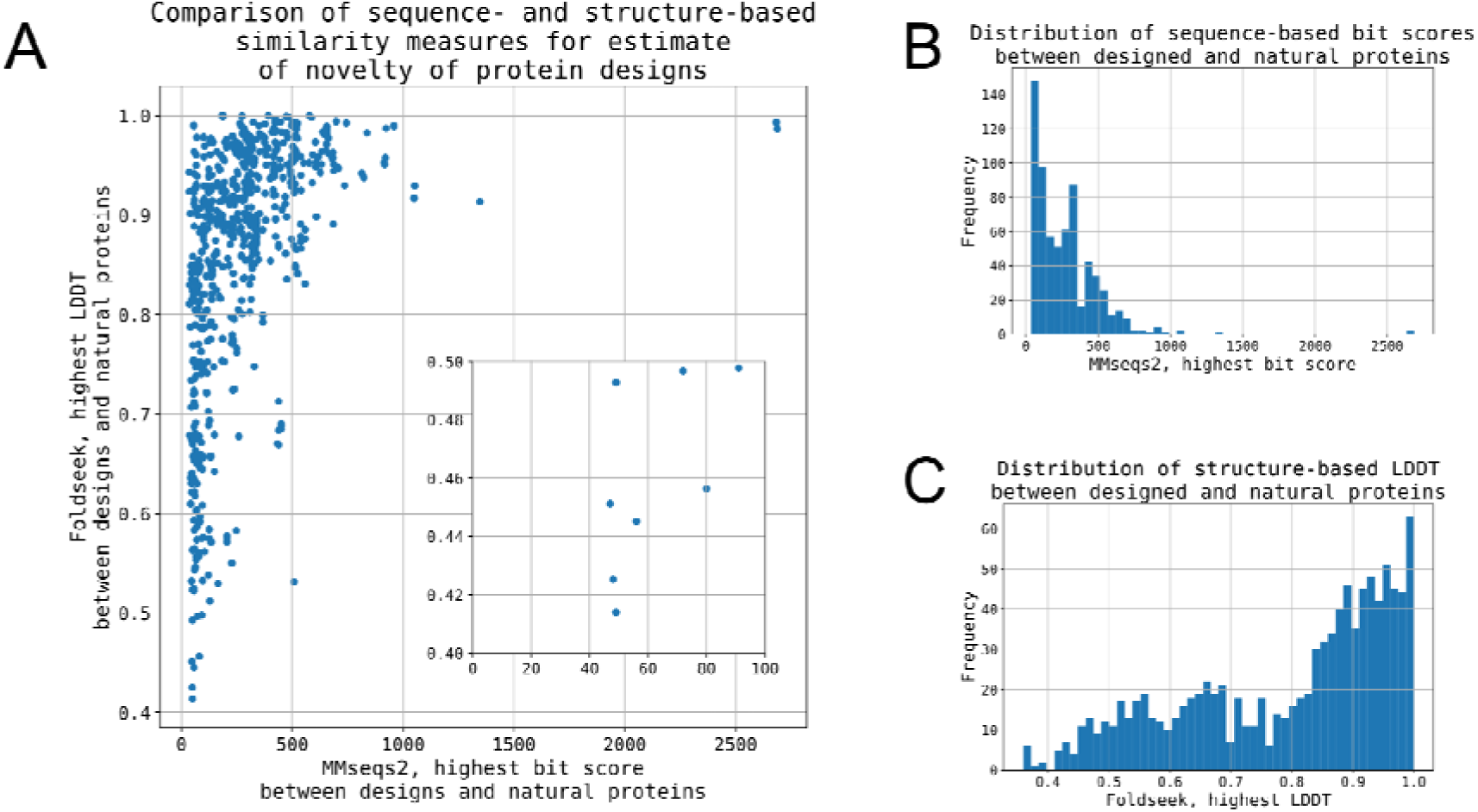
A) Correlation between the highest sequence-based bit scores and structure-based LDDT for every PDA entry, compared to natural proteins with an earlier release date than query (DvP-type). In the bit score range below 1500, data shows 50% Pearson’s correlation coefficient. B) Distribution of sequence-based bit scores, the highest values for every entry within the PDA dataset, compared to natural proteins with an earlier release date than query (DvP-type). C) Distribution of structure-based LDDT, the highest values for every entry within the PDA dataset, compared to natural proteins with an earlier release date than query (DvP-type).

We have found that 139 designs have no sequence similarity, and 268 designs have no structural similarity to valid natural targets. These, excluding designs that did not produce output due to a range of factors such as being too short, account for 17% and 21% of tested designs, respectively. Full list of designs with no natural matches is reported in Supplementary Information table S3 and S4.

Other notable examples from the analysis include 1qys (Top7), which is shown to have a distinct, but structurally related fold to 1lfw, a bacterial protein, while showing no sequence similarity to any known natural target (fig. S3A). The two designs which appear to have extremely high similarity (bit scores > 2250 and LDDT > 99%) to natural targets, 8a1a and 8a19, are in fact valid designs, with designed sequences fused with those of natural origin (DARPin–leucinostatin derivatives, fig. S3B). These examples highlight the challenges in working with biological metadata.

For any two different designed chains (DvD-type), the lowest sequence identity match is 20.5%, and the lowest bit score was 27; for designed chain and any of the natural sequences in the PDB (DvP-type) it was 18.7% and 35, respectively. 86% of the PDA entries exhibit 50% or higher sequence identity to natural proteins, and 95% have bit scores equal to 50 or higher. This suggests that protein designs are still inspired by natural structures, and there continues to be difficulty in designing entirely novel structures ^64,65^.

We report the highest scoring protein partners (both designed and natural) on each entry’s details page, which should not only map interesting relationships in the data but also reveal “orphaned” designs that have not been published in scientific journals, yet still deposited in the PDB.

## Discussion

This paper presents the Protein Design Archive (PDA) database. This is an up-to-date resource containing all *de novo* protein designs, which is readily searchable and can be analysed to gain insights into the field of protein design. As it has been repeatedly claimed that this research area has “come of age” ^12^, and it has delivered successful achievements such as small molecule binders ^66,67^, protein-protein binders ^68,69^, and entirely novel folds touching the “dark matter” of protein space ^41,47,70^, there is a huge amount of information that we can learn from existing designed proteins, that could be used to inform and improve design methods in the future.

Over the past 40 years of protein design, the field has produced around 50 structures a year, but developments in computational methods and software accessibility have hugely aided this process, speeding the rate of release of new designs almost 10-fold, and prompting us to divide the growth curve of number of protein designs into two distinct periods (fig. 2). In recent years (2020 – 2023), the number of new designed structures deposited in the RCSB PDB database has doubled compared to the decade before that (2010 – 2020). This might be attributable to the rise of deep learning-based approaches in protein design.

We have found that in general, but especially in the early days of protein design, there tends to be a lack of set rules for labelling data, with overlap between phrases and labels that protein designers and engineers use. Some entries are also simply difficult to define, due to no quantifiable and universally accepted definition of what is *de novo* designed and what is engineered. With the PDA, we have been as inclusive as possible with our definition of designed proteins and have given the user tools to restrict the proteins in the archive according to their own definition, if they disagree with ours. Furthermore, we have identified currently “orphaned” designs (161 excluding releases in the last 12 months, 233 in total), which were previously hard to find as they have not been described in the published literature.

Another challenge in biological metadata has been posed by labelling of designed chains. In our analysis, we aimed to include only designed chains, based on the chain source labels in the CIF files. However, we have encountered discrepancies between chain labels between CIF, FASTA, and PDB files. While FASTA files utilised by MMseqs2 conveniently carry the chain source information within their description, PDB files do not. With this in mind, we are aware that selection of designed chains for the Foldseek analysis could be improved, although the analysis still provides useful insights. This issue highlights both the need for better structuring of biological metadata, as well as the usefulness of the PDA database.

While other broad studies of designed proteins exist ^7,9,12,55–59^, none of them have aimed to systematically capture all protein designs. By necessity, their curation was performed by hand, and could not be continued easily post-publication. Furthermore, often, they do not include detailed methodology of how the curation was performed, making it difficult to extend their analyses. Those that do, reveal much smaller, closer to 100-entries-large collections, suggesting that some designs have been omitted. The PDA database addresses many of these issues, with the data readily available for download on our website and the methods described in detail to ensure reproducibility.

Key findings from our analysis show that designed proteins have a bias towards specific amino-acid usage and secondary structure types when compared to the PDB as a whole. Designed proteins have a larger proportion of α-helical secondary structure than proteins in the wider PDB. Despite these differences, there is evidence that designed proteins are becoming closer to natural proteins in the PDB in terms of the complexity of the proteins being designed, with a larger range of amino acids being incorporated into designs in more recent years, and the size of the designs increasing over time. At the same time, it remains a goal of *de novo* protein designs to create novel sequences and structures, exploring possibilities outside nature. Our analysis highlights achievements worth celebrating, while reflecting the overall modest success.

Historically, structural data has been critical to the development and validation of protein design methods, with X-ray crystallography providing the accurate 3D data required for validating the design, and often revealing opportunities for further optimization. The growing trend of utilising cryo-EM to reveal larger and more-complex structures and assemblies highlights its increasing importance in structural biology and its ability to tackle increasingly complex designed protein structures.

However, structural data only accounts for a tiny amount of the information collected regarding designed proteins and protein design methodologies, and these data only really represent “successful” designs. If the field is to progress, information regarding all aspects of the design process must be collected. It is our intention that the PDA will be expanded to include information on design protocols, sequence information, as well as results of biochemical and biophysical analysis of designs in the future.

## Methods

### Data collection

The data was collected through automated querying of the RCSB PDB ^71^, with the search query focused on the mmCIF dictionary definition of polypeptide (*_entity_poly.type*) derived from synthetic constructs, as defined by *entity_src_syn.ncbi_taxonomy_id* (32630) ^72^. This information allowed identification of the original mmCIF dataset of *de novo* designs, which was then further processed. The *_entity.pdbx_description* field within this dataset provides the mmCIF definition corresponding to the macromolecule’s name provided by the authors at time of deposition.

The results of the automated query of the RCSB PDB database were curated by the paper’s authors. To finally complete the dataset to our best ability, we supported the automated search by manual inspection of the RCSB PDB database based on authors and keywords, literature review, and knowledge of the field.

The database is updated monthly by its authors running the automatic search query and manually reviewing the designs to maintain high standard of included data. The date of the most recent update is displayed at the bottom of each page, along with the number of protein designs added.

The authors eagerly encourage protein designers to contact them about missing information and feedback.

### Data processing

For each design we collected detailed structural and sequence information (below, fields starting with an underscore symbol refer to their label in the CIF file):

- PDB code
- release date (_pdbx_audit_revision_history.revision_date)
- subtitle (_struct.title)
- tags (_struct_keywords.text and _struct_keywords.pdbx_keywords)
- the full publication abstract, and the keywords extracted from it using the Natural Language Toolkit for Natural Language Processing with Python ^73^
- authors (_citation_author.name if _citation_author.citation_id is primary)
- publication citation (including _citation.title, _citation.journal_abbrev, _citation.journal_volume, _citation.page_first, and _citation.page_last), publication link (_citation.pdbx_database_id_{DOI / PubMed / CSD / ISSN / ASTM}), and publication country (_citation.country)
- for each chain (_entity_poly.entity_id):

- CIF file-based chain IDs (_entity_poly.pdbx_strand_id), primary FASTA file-based chain IDs (comma separated within chain description), and author-assigned FASTA file-based chain IDs (inside “[auth]” brackets, within chain description)
- Chain source, described in the

_entity_src_gen.pdbx_gene_src_scientific_name or
_pdbx_entity_src_syn.organism_scientific field, depending on whether the molecular entity label matched the _entity_src_gen.entity_id or
_pdbx_entity_src_syn.entity_id fields, respectively
- Chain type based on the source organism:

▪ “D” for “designed” if chain source fields contain phrases "synthetic construct" or "artificial",
▪ “U” for “unknown” if chain source fields are missing, or contain phrases “unknown”, “unidentified”, or “?”,
▪ Otherwise “N” for “natural”, unless the design entry would then seem to contain only natural chains, then “M” for “missing”,
- amino acid sequences with indications of unnatural amino acids (_entity_poly.pdbx_seq_one_letter_code) and with natural amino acid equivalents (_entity_poly.pdbx_seq_one_letter_code_can)
- sequence length (sum of letters in _entity_poly.pdbx_seq_one_letter_code)
- static 2D image of the biological assembly, and a 3D interactive structure using NGL viewer ^60,61^
- crystal structure information (_cell.length_{a / b / c}, _cell.angle_{alpha / beta / gamma})
- symmetry (_symmetry.space_group_name_H-M)
- formula weight (_entity.formula_weight)
- experimental method of characterisation (_pdbx_entity_src_syn.details)
- synthesis comment (_pdbx_entity_src_syn.details)

We did not make the PDA a calque of the RCSB PDB, instead removing redundant fields, to make the database interesting, perspicuous, and easy to browse. For further needs, we refer users to the RCSB PDB webpage through hyperlinks to the relevant data entries.

### Analysis of Designed Proteins vs the PDB

The PDB files of 1,432 designed proteins in the PDA database were downloaded from the Protein Data Bank (PDB) ^71^ and only the chains of the designed protein were kept (Supplementary Information fig. 4). All other chains in the PDB files containing proteins that were not designed, were stripped out of the PDB files. After this, the amino acid proportions were calculated for these designed proteins, with the non-standard residues being converted to the closest standard amino acid or being classed as non-standard residues (‘X’). In addition to this, DSSP ^74^ secondary structure proportions were calculated along with other metrics capturing all-atom scores, geometric properties, and aggregation propensities from the DE-STRESS software. The designs were then grouped by their release year into 5-year intervals and all these metrics were averaged across these groups. Furthermore, this analysis was repeated for 216,681 proteins from the PDB (up to date as of June 2024) which was used as a comparison for the designed proteins.

MMseq2 and Foldseek “easy-search” modules were ran with the highest possible sensitivity (-s 7.5) and iteratively (--exhaustive-search). Two types of analysis were executed: DvP-type search, where designed proteins were compared against natural targets, and DvD-type search, where designs were compared against themselves.

Query and target databases were built using “createdb” module (219,821 proteins from the PDB, up to date as of August 2024), with the exception for Foldseek’s PDB target database, where a pre-generated PDB database was used, downloaded using Foldseek’s “databases” module.

The analysis included single, and only designed chains. For MMseqs2, designed chains were selected and included in the analysis based on the FASTA file description fields. Chains which contain “32630” (taxonomy id for synthetic construct) in source were kept. For Foldseek, chains were labelled as “designed” and included in the analysis based on matching their chain labels in the PDB file with labels given in CIF file (labelling method described in “Data processing” section), or primary labels in FASTA file, or author-assigned labels in FASTA file. First, CIF files were utilized to identify chain labels and determine which chains were designed. Then, chain labels were scraped from FASTA files, where the chain descriptions sometimes differed from those in the CIF files. To reconcile these differences, FASTA chain labels (both primary and author-assigned) were matched with those from the CIF dataset based on shared PDB codes. Then, it was assumed that each PDB file consistently uses a single chain labelling system, but different PDB files might use different systems. To account for this, chain labels from the PDB file were checked against designed chains, sequentially matching CIF file labels, or primary labels in FASTA file, or author-assigned labels in FASTA file, including matching chains into the analysis.

For MMseqs2, metrics measured were: bit scores (“bits”) and percentage of identical matches (“pident”). For Foldseek, these were: probability of being homologous (“prob”), average local distance difference test (“lddt”), alignment TM-score “alntmscore”, e-value (“eval”), and bit scores (“bits”) ^75^.

When reporting the similarity results, hits between query and target were excluded when query and target are from the same protein. Additionally, for DvP-type analysis, hits were excluded when target is another designed protein, or target has been deposited to the RCSB PDB later than the query.

Details of the number of inputs for software, outputs of software, and outputs of analysis (as unique PDB codes and unique chains) are summarised in the Supplementary Information table S3. For structure-based search, only 1432 PDA database entries were analysed, due to the PDB format files being not available for large structures. The number of chains selected as “designed” for sequence- and structure-based analysis differs, due to unavailability of PDB files of large structures, and due to a different method of selecting chains (described above). Input entries may be missing from software output for a range of reasons, including queries being too short to be aligned with targets: e.g. MMseqs2 requires a minimum sequence length of 14 residues to make a match. Software output entries missing from analysis results are significant, as they are the designs that have not been found to have any sequence or structure similarity with any natural and older protein. Full list of these is included in the Supplementary Information table S4.

## Data availability

All data generated in this study is available on the PDA Database website, downloadable as JSON or CSV file, and on GitHub, at the following address: https://github.com/wells-wood-research/prot-des-timeline.

## Code availability

The scripts for website build, data collection and processing, and protein designs analysis used in this study are publicly available on GitHub, at the following address: https://github.com/wells-wood-research/prot-des-timeline.

## Acknowledgements

We thank members of the Wells Wood Research Group for testing and feedback on the PDA website. Marta Chronowska is supported by a PhD studentship from the UKRI funded EastBio Doctoral Training Partnership programme. Michael J. Stam, Christopher W. Wood, and Derek N. Woolfson are supported by a BBSRC sLOLA award (BB/X003027/1).

