## Supplementary Information for "The Protein Design Archive (PDA): insights from 40 years of protein design"

### The Protein Design Archive Supplementary Information

#### Growth of protein designs over time:

| **statistic** | **Region 1 (minimal and rational)** | **Region 2 (computational)** |
| --- | --- | --- |
| **Slope** | 10.131868131868131 | 88.74175824175825 |
| **Intercept** | -20233.76923076923 | -178366.8956043956 |
| **R-squared** | 0.9729425100443526 | 0.9696795574539044 |
| **p-value** | 8.956418746890206e-11 | 1.0631114981979551e-09 |
| **Standard error** | 0.48775201149273767 | 4.731352815566564 |

Table S1: Statistics of the linear fits of the protein design growth curve (fig. 2), separated into two regions based on the dominant approaches to protein design over these time periods *minimal and rational (1997 – 2010), and computational (from 2010).*

| **year** | **count** |
| --- | --- |
| 1991 | 1 |
| 1992 | 1 |
| 1993 | 1 |
| 1997 | 4 |
| 1998 | 7 |
| 1999 | 4 |
| 2000 | 4 |
| 2001 | 12 |
| 2002 | 12 |
| 2003 | 15 |
| 2004 | 18 |
| 2005 | 7 |
| 2006 | 4 |
| 2007 | 5 |
| 2008 | 9 |
| 2009 | 10 |
| 2010 | 32 |
| 2011 | 25 |
| 2012 | 70 |
| 2013 | 26 |
| 2014 | 74 |
| 2015 | 76 |
| 2016 | 85 |
| 2017 | 83 |
| 2018 | 80 |
| 2019 | 58 |
| 2020 | 140 |
| 2021 | 92 |
| 2022 | 144 |
| 2023 | 200 |
| 2024 | 151 |

Table S2. Number of new protein designs released by year.

#### Similarity analysis:

##### Structure-based:

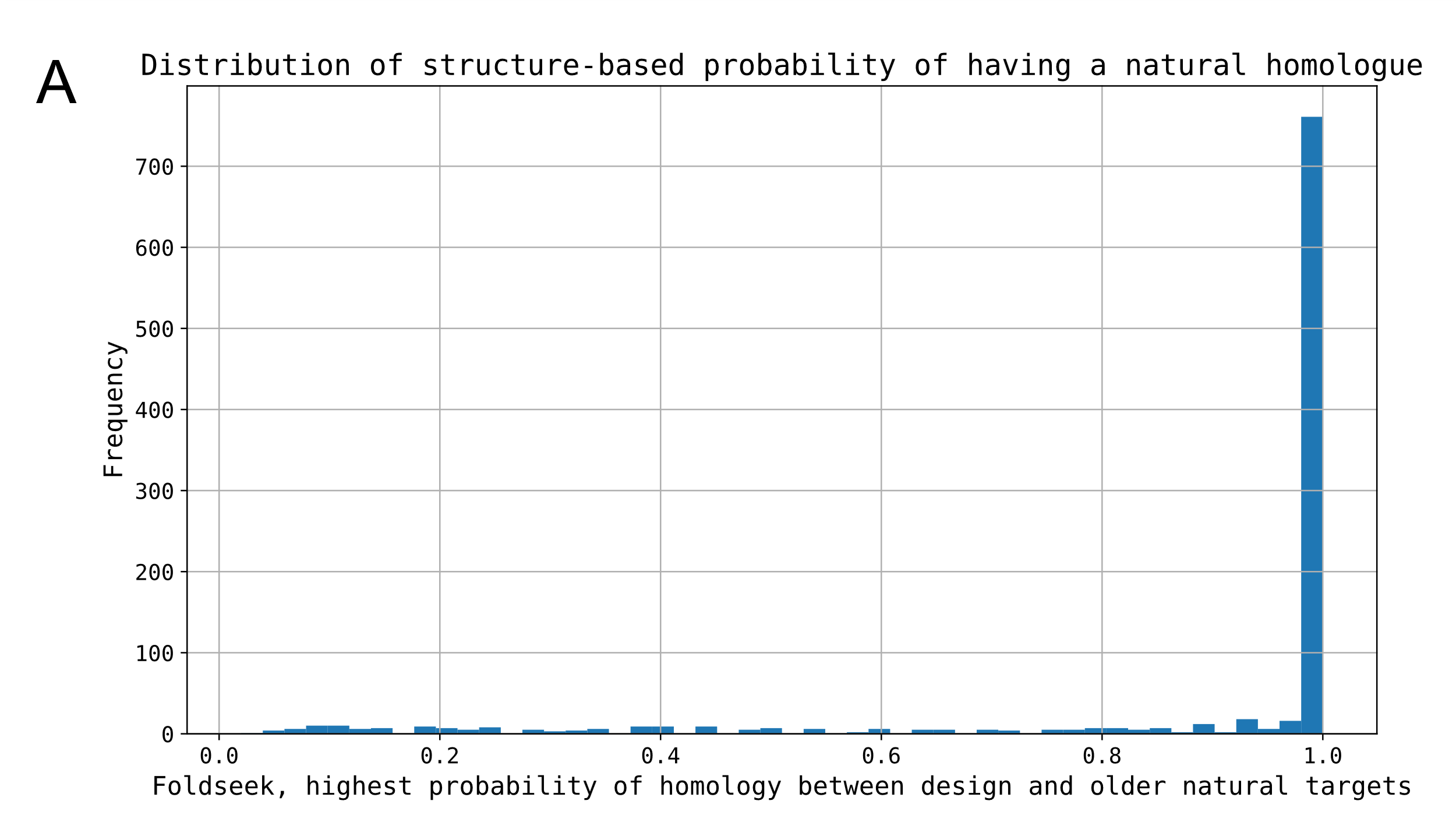

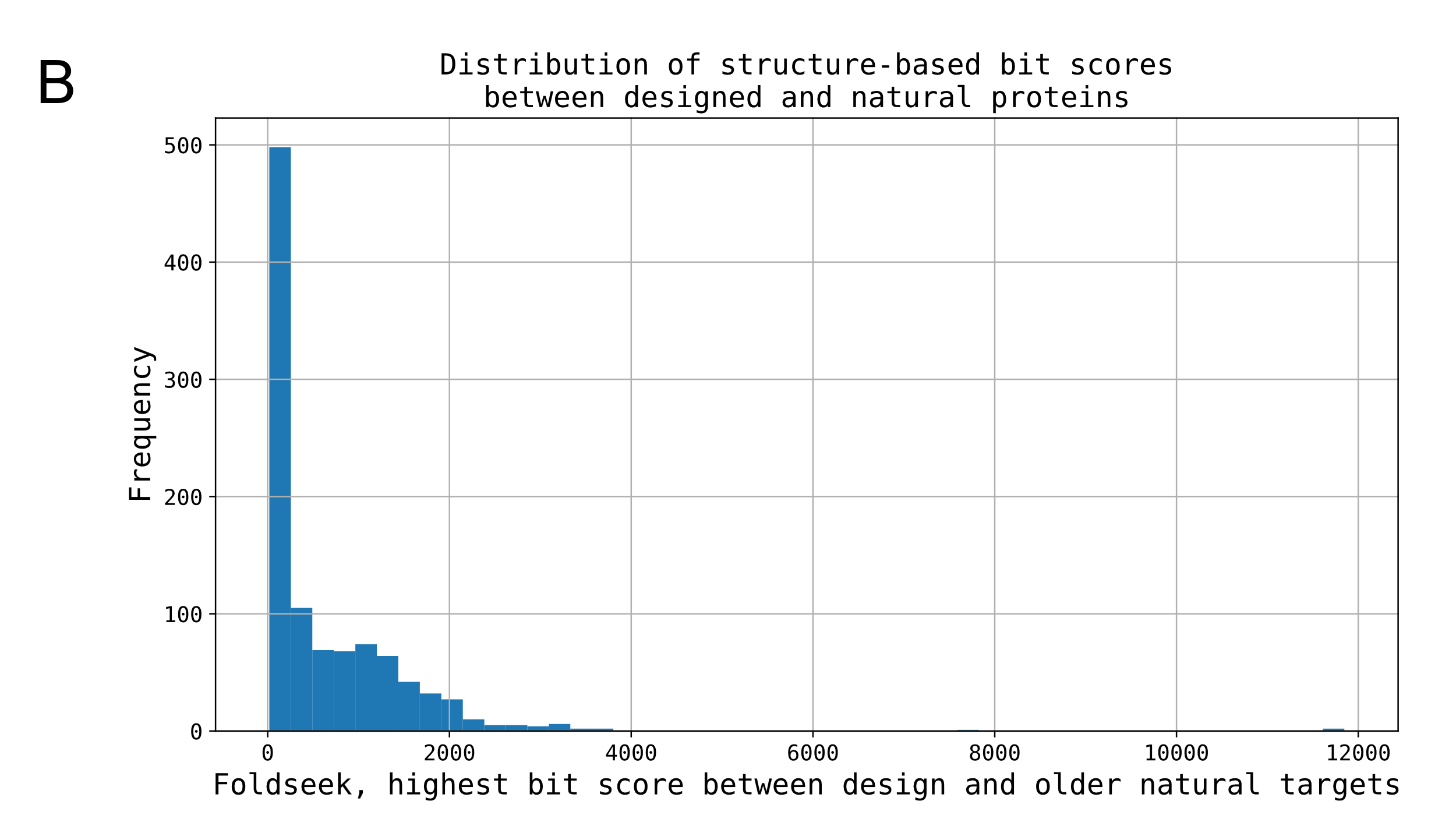

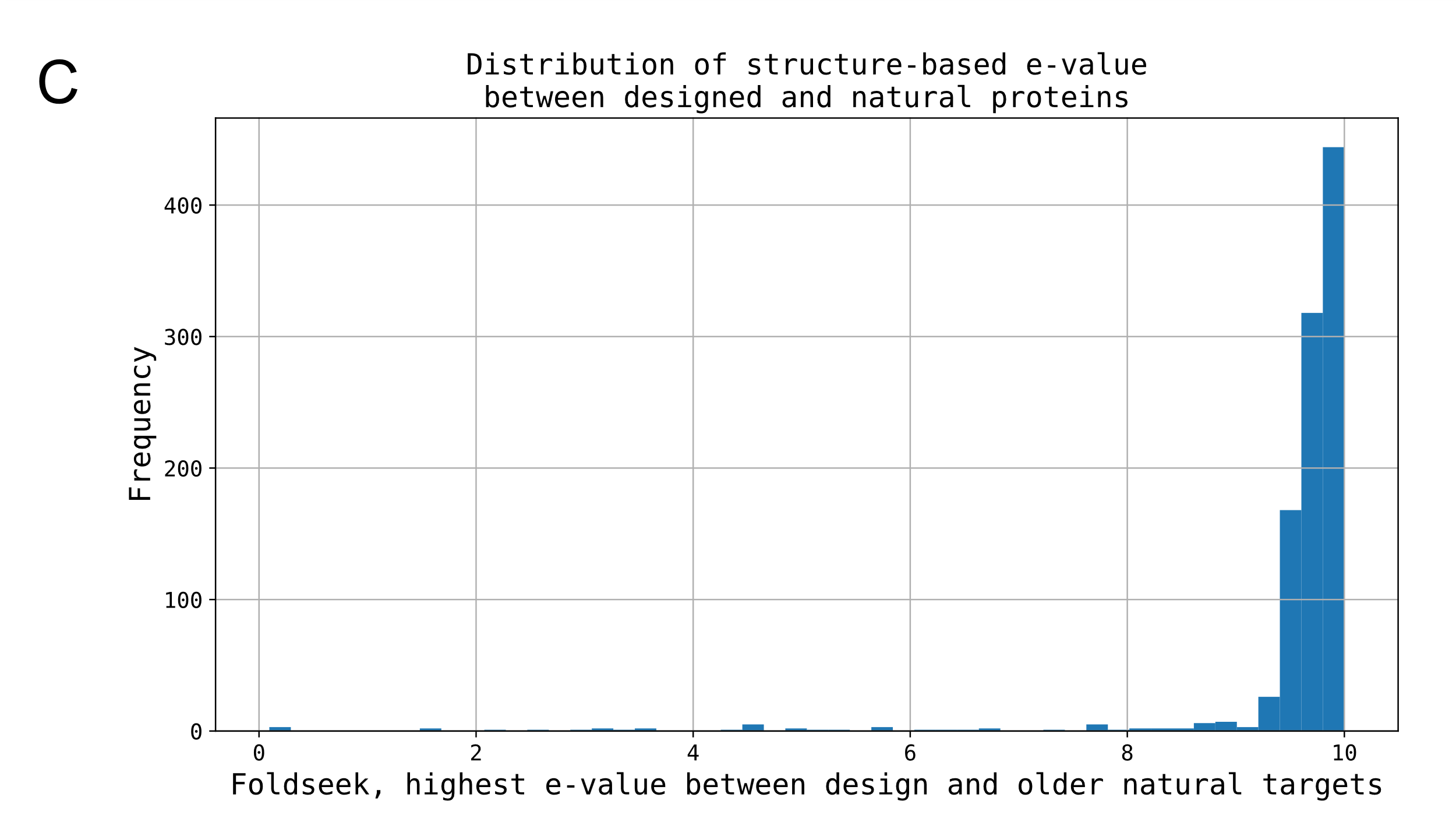

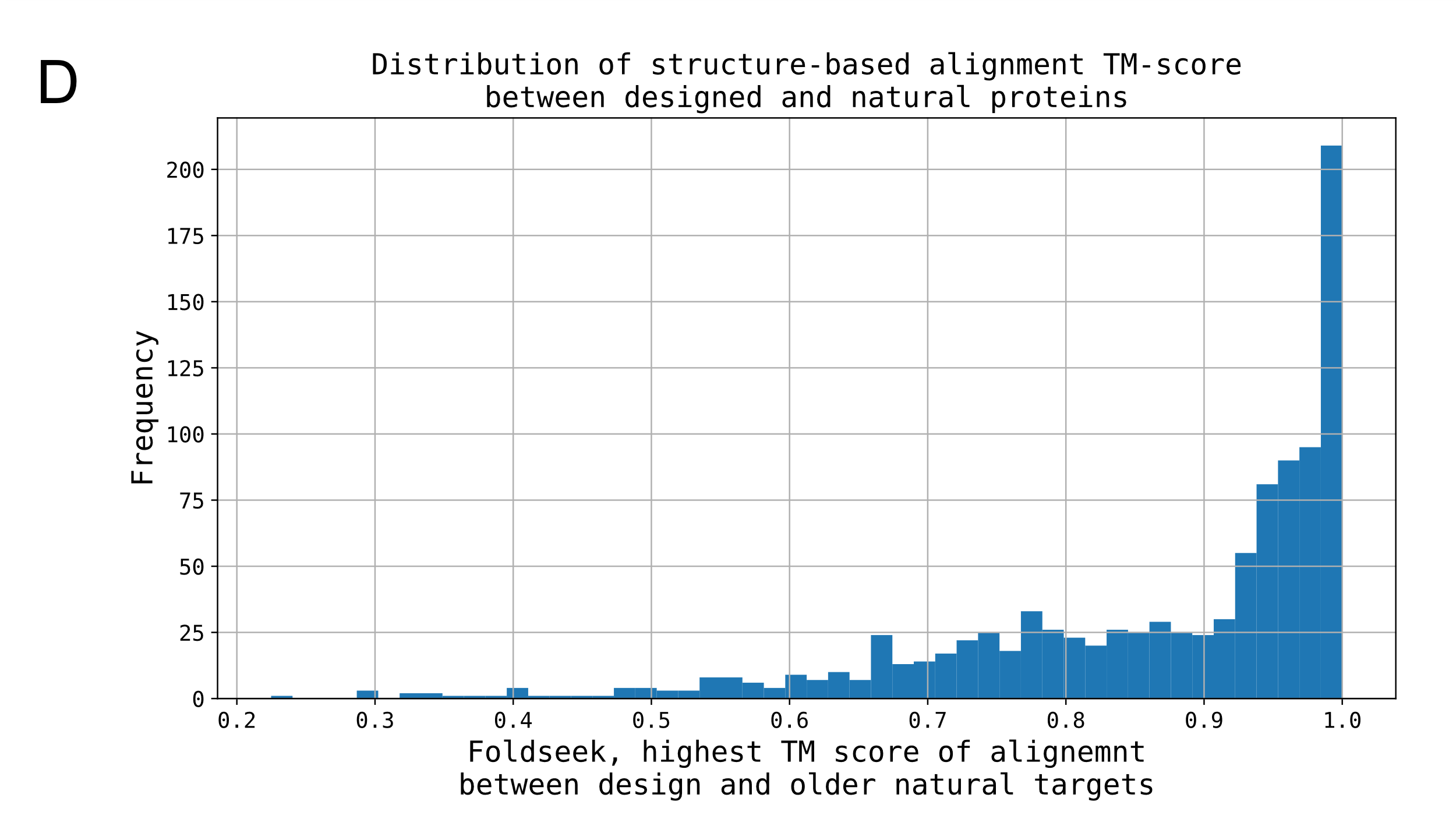

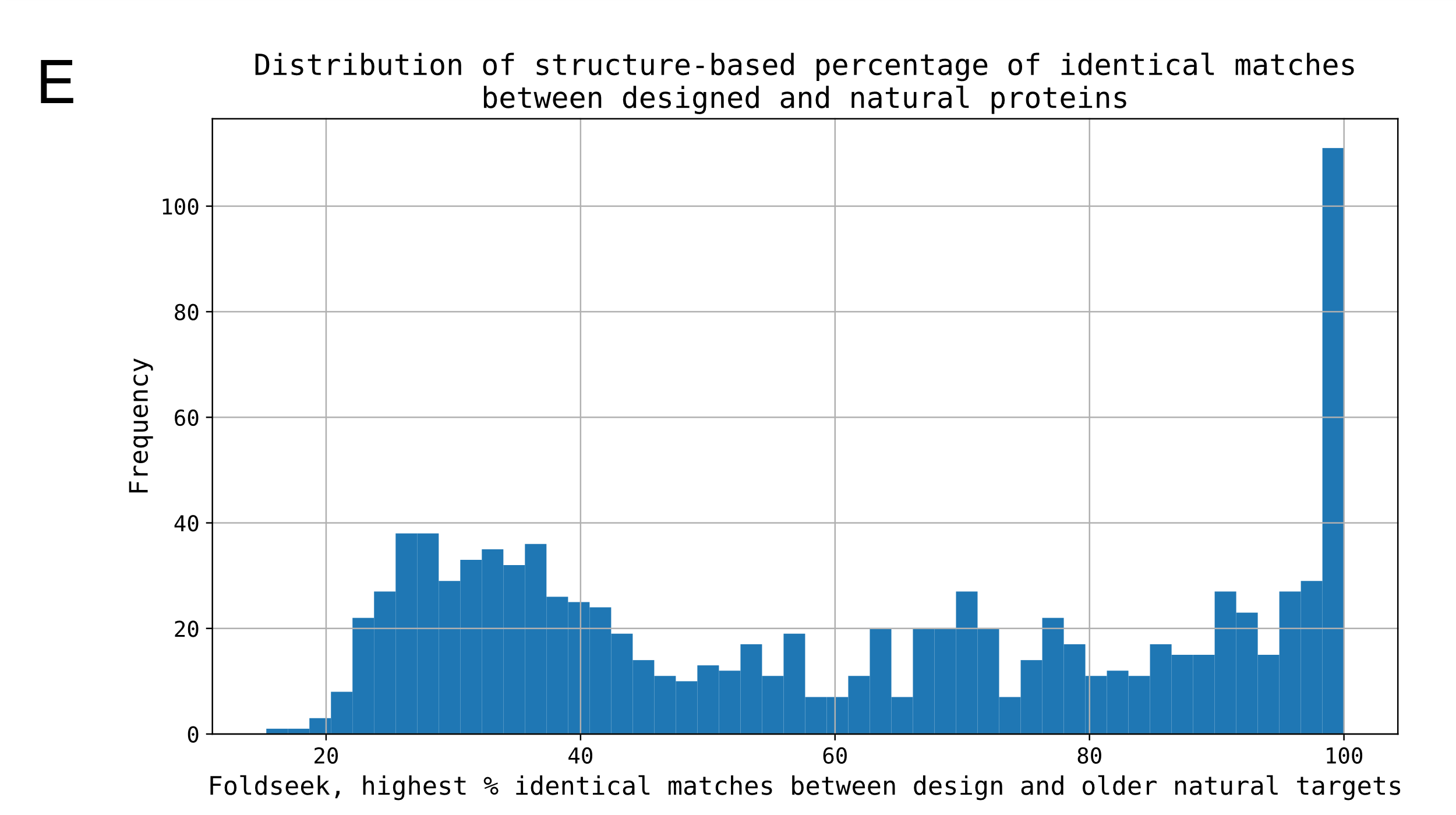

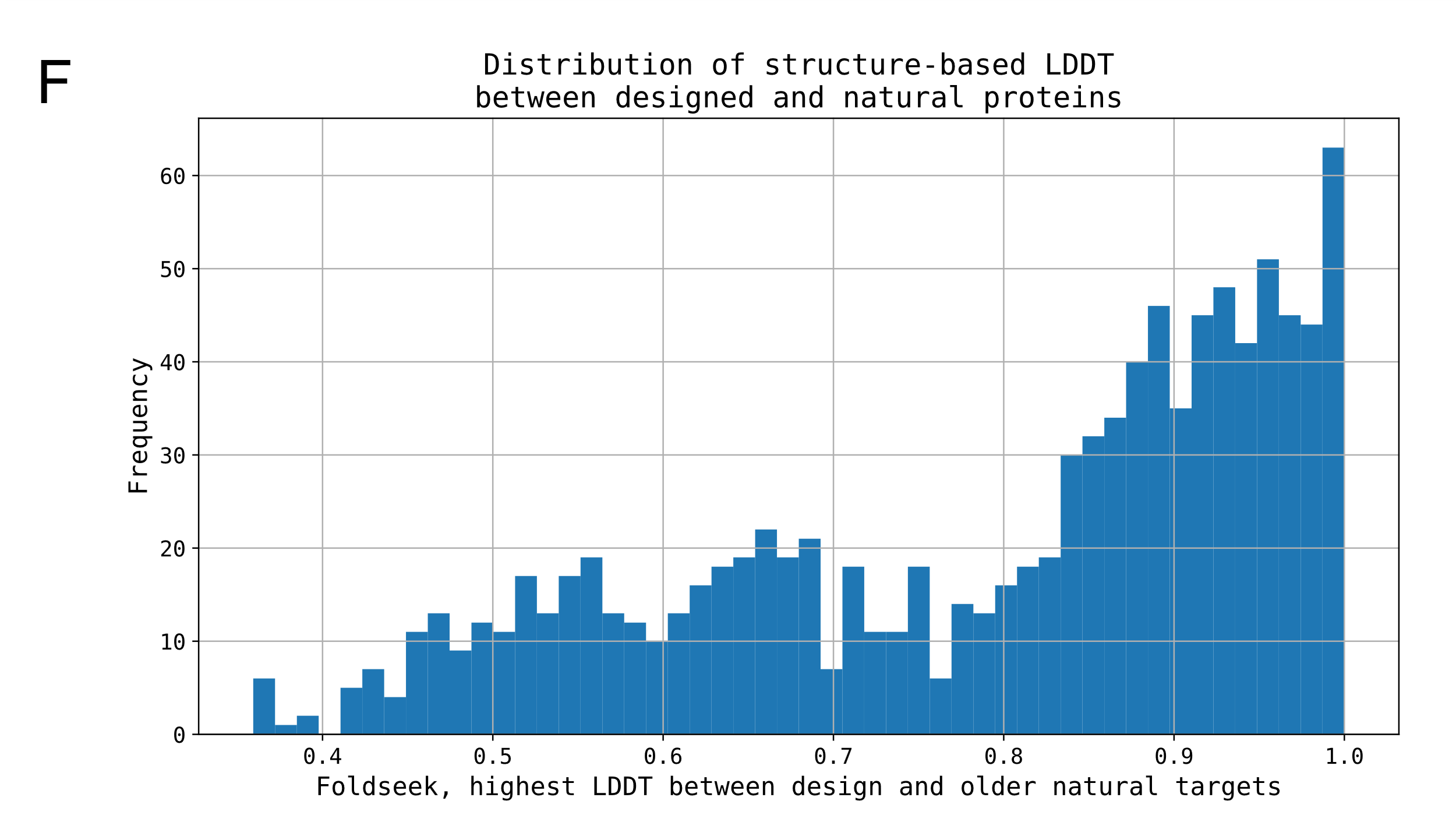

Figure S1: Distribution of distribution of various Foldseek (structure)-based similarity metrics: A) probability of being homologous, B) bit scores, C) e-value, D) TM score of alignment, E) percentage of identical matches, F) LDDT. For each metric and for every design, the highest value scored between the designed query and natural protein target was recorded.

##### Sequence-based:

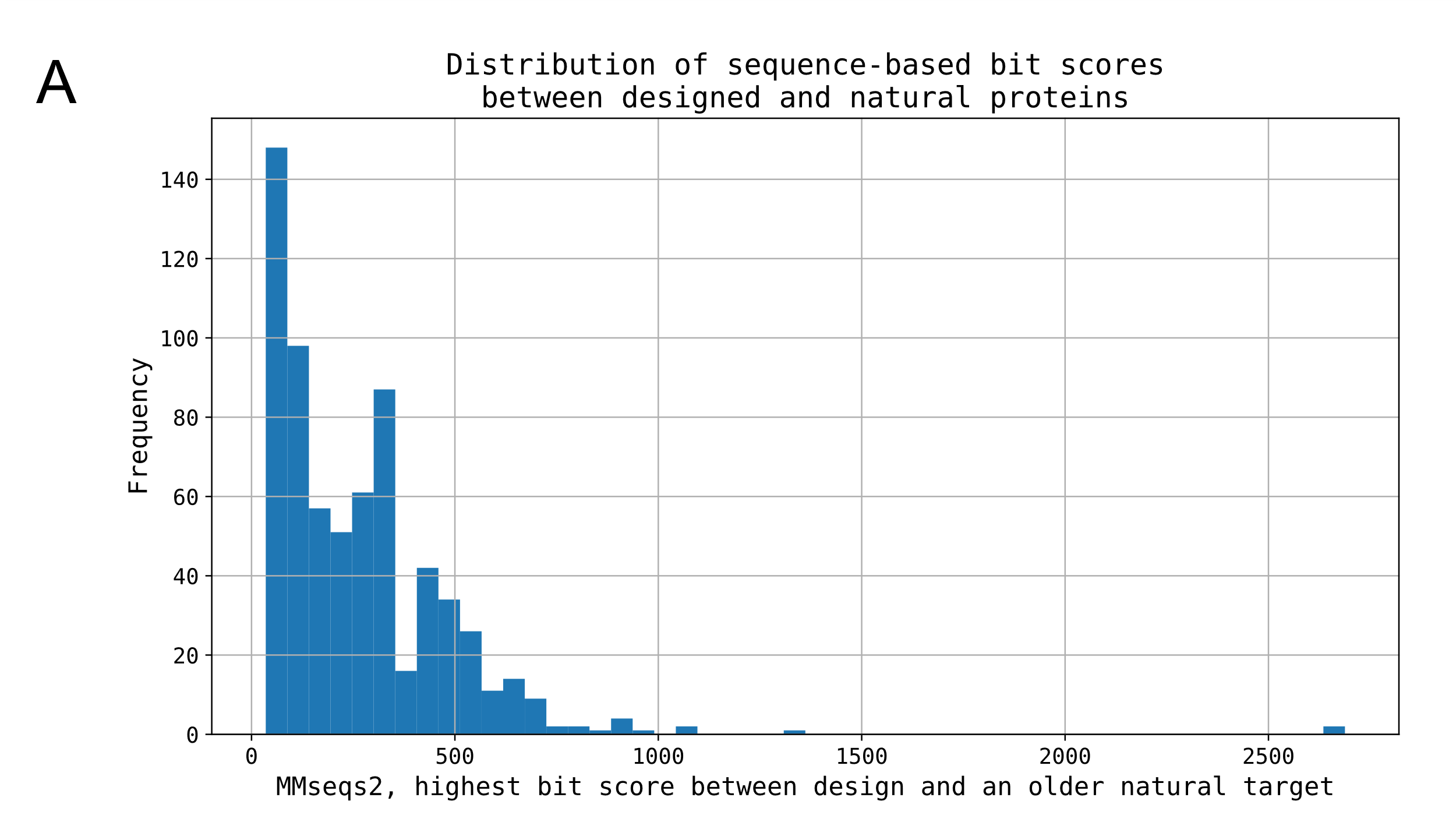

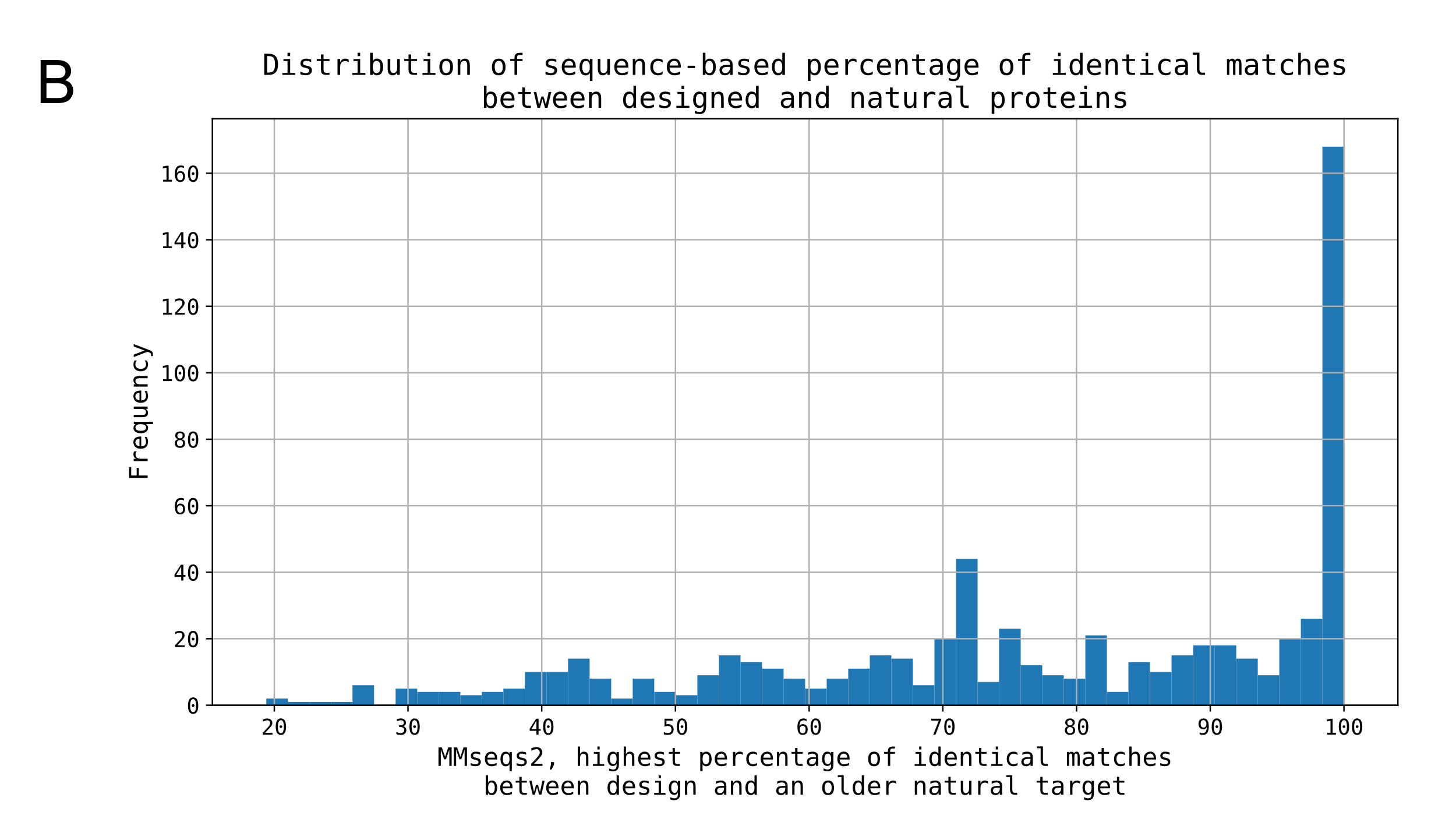

Figure S2: Distribution of distribution of various MMseqs2 (sequence)-based similarity metrics: A) bit scores, B) percentage of identical matches. For each metric and for every design, the highest value scored between the designed query and natural protein target was recorded.

##### Notable examples:

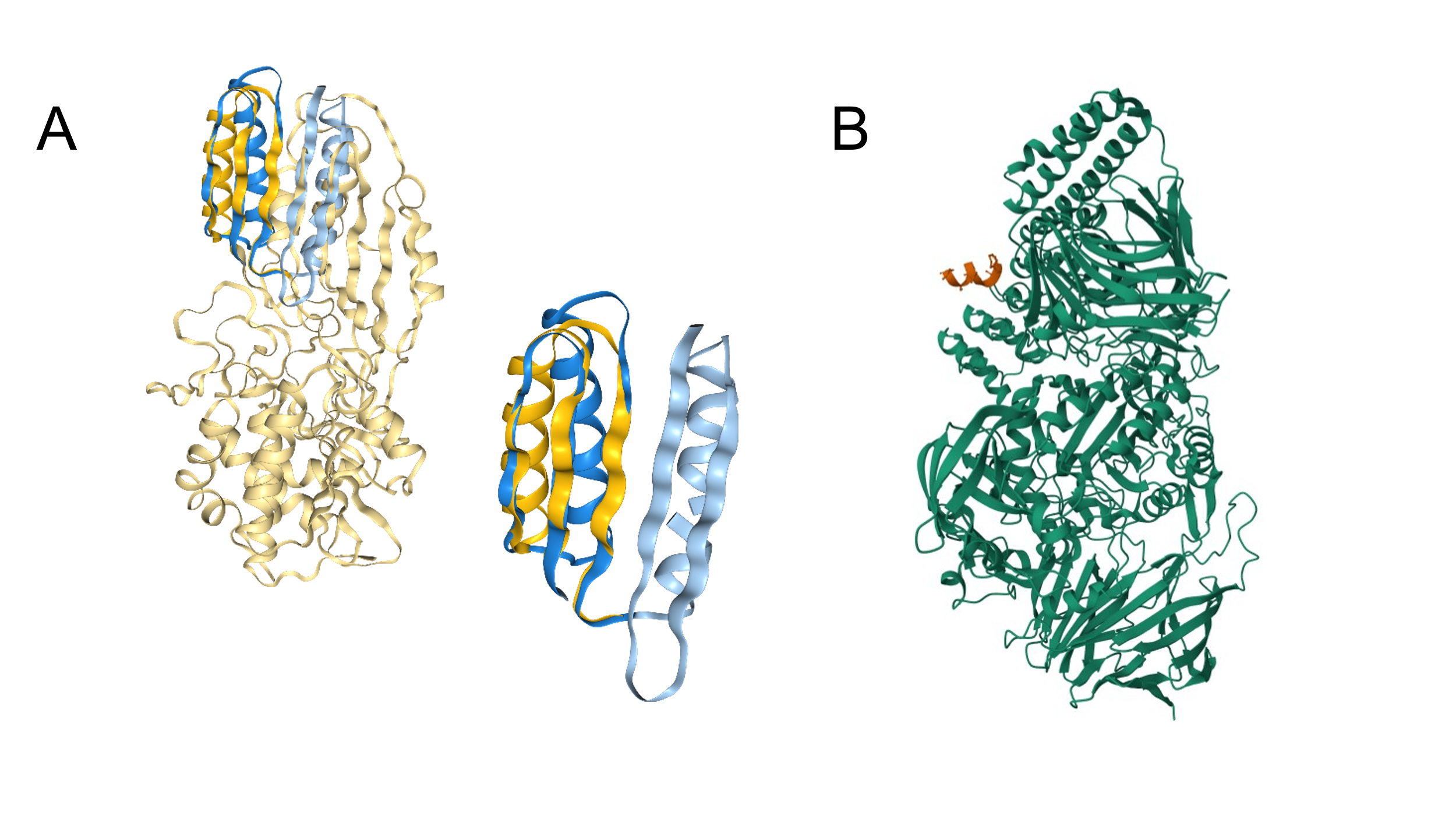

Figure S3: Notable examples from the analysis: A) fold of de novo design 1qys (Top7) is distinct but structurally related to 1lfw, a bacterial protein, while showing no sequence similarity to any known natural target. B) Two designs which appear to have extremely high similarity (bit scores > 2250 and LDDT > 99%) to natural targets, 8a1a and 8a19, are in fact valid designs, with designed sequences fused with those of natural origin (DARPin–leucinostatin derivatives).

##### Comparison of sequence/structure similarity correlation for concatenated chains vs single chains:

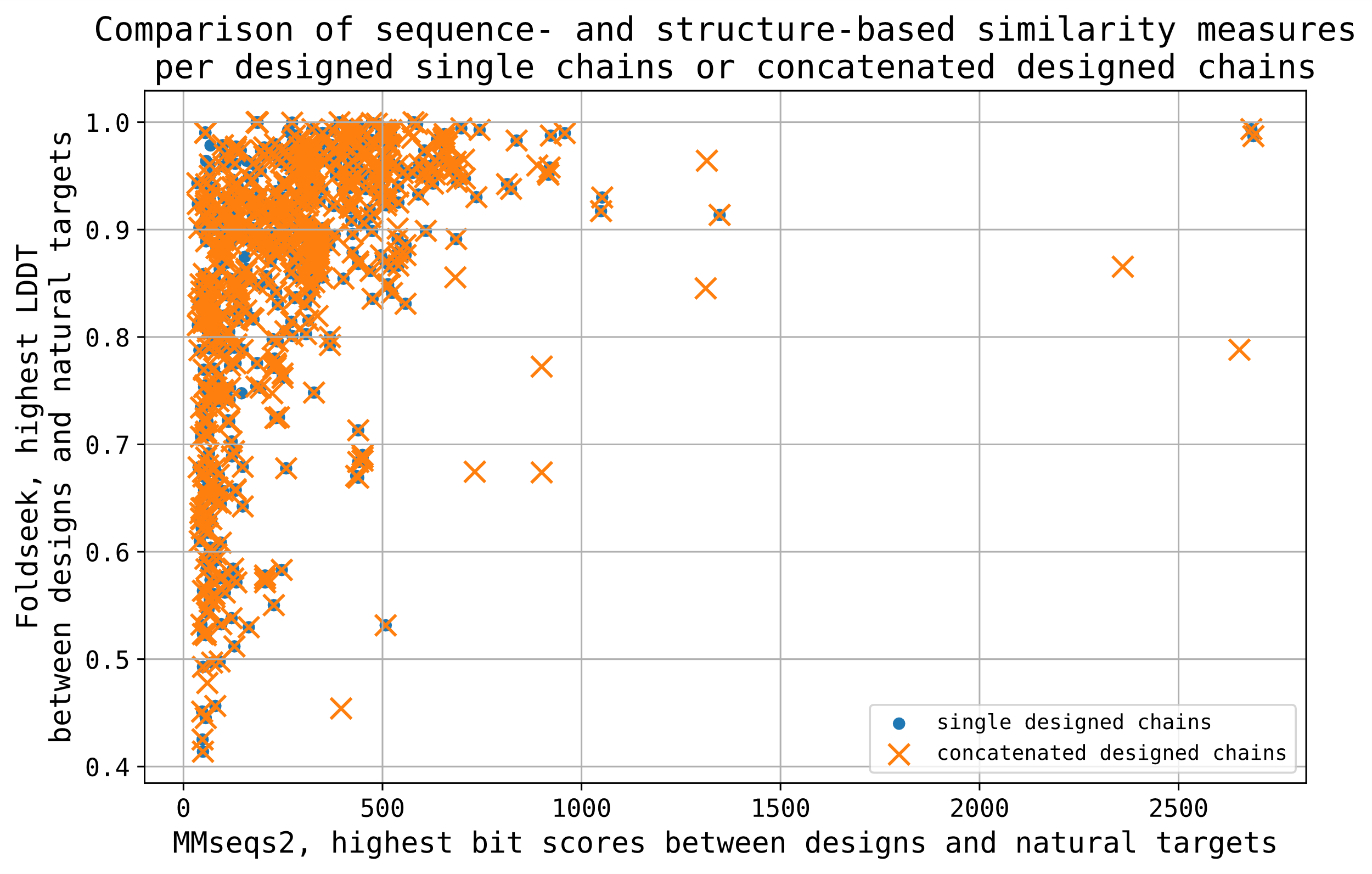

Figure S4: Comparison of correlation between sequence-based bit scores and structure-based LDDT when measured for per single designed chains or concatenated designed chains. With three exceptions, the results are the same. We proceeded with single chain-based analysis throughout this paper.

##### Number of entries passing through the analysis pipeline contains important information:

| **Software** | **Type – design vs design (DvD) or design vs natural protein (DvP)** | **Number of inputs for software** | | **Number of outputs of software ^D)^** | | **Output of analysis ^E)^** |
| --- | --- | --- | --- | --- | --- | --- |
|  |  | **# unique PDB codes** | **# unique chains** | **# unique PDB codes** | **# unique chains** | **# unique PDB codes** |
| **MMseqs2** | DvD | 1450 | 1592 ^B)^ | 1167  (1450 ^F)^) | 1268  (1592 ^F)^) | 932 |
|  | DvP | 1450 | 1592 | 808 | 869 | 669 |
| **Foldseek** | DvD | 1432 ^A)^ | 1603 ^C)^ | 1350 | 1374 | 1270 |
|  | DvP | 1432 | 1603 | 1284 | 1307 | 1016 |
| **DE-STRESS** | - | 1432 | - | 1247 ^G)^ | - | - |

Table S3: Summary of the number of entries (by unique PDB codes and unique single chains) at every step of the analysis: software (MMseqs2, Foldseek, DE-STRESS) inputs, software outputs, and analysis outputs. The number of entries that passed every step is significant and informs of many nuances to the analysis. A) PDB format files are not available for large structures, hence not all PDA entries were analysed by Foldseek and DE-STRESS. B) For MMSeqs2, chains were labelled as “designed” and included in the analysis based on the FASTA file description fields. Chains which contain “32630” (taxonomy id for synthetic construct) in source were kept. C) For Foldseek, method of selection of designed chains was limited by biological metadata being inconsistent between different file formats. This limitation resulted in number of designed chains included in Foldseek and DE-STRESS analysis being greater than the number analysed using MMseqs2, known to label designed chains much more reliably. D) Input entries may be missing from software output for a range of reasons, including queries being too short to be aligned with targets, or query not finding any match besides itself. E) Software output entries missing from analysis results are significant, as they are the designs that have not been found to have any sequence/structure similarity with any natural and older protein. F) To test how many designed queries can pass the software analysis, flag “--add-self-matches 1” was used in the DvD-type analysis. This ensured that the input entries are not missing from software output due to errors, but only due to not finding a hit with a target different than itself. When the flag including in output matches of query with itself was used, all 1450 designed pdb entries and 1592 designed chains passed the software analysis successfully. This shows that all input sequences are able to pass the analysis, and entries missing from DvP-type search are due to not finding matches with anything besides themselves, supporting their originality as *de novo* protein designs.

| **Steps of analysis being compared** | **PDB codes of entries missing between the two analysis steps** |
| --- | --- |
| MMseqs2 input and analysis results | 1abz, 1al1, 1bb1, 1byz, 1coi, 1cos, 1d7t, 1djf, 1ec5, 1fme, 1fmh, 1fsd, 1fsv, 1g6u, 1hcw, 1hqj, 1ic9, 1icl, 1ico, 1j4m, 1jm0, 1jmb, 1jy4, 1jy6, 1jy9, 1k43, 1kd8, 1kd9, 1kdd, 1kyc, 1l4x, 1le0, 1le1, 1le3, 1lq7, 1m3w, 1n09, 1n0a, 1n0c, 1n0d, 1nvo, 1p68, 1pbz, 1psv, 1pyz, 1qp6, 1qys, 1rh4, 1s9z, 1sn9, 1sna, 1sne, 1t8j, 1tgg, 1tjb, 1u0i, 1u2u, 1uno, 1uw1, 1vl3, 1vm4, 1vm5, 1vrz, 1xof, 2a3d, 2cw1, 2evq, 2gjh, 2jgo, 2jua, 2jvf, 2k6r, 2ki0, 2kjn, 2kjo, 2kl8, 2klw, 2koz, 2kp0, 2kpo, 2l69, 2l82, 2l96, 2l99, 2l9a, 2lci, 2ln3, 2lnd, 2lny, 2lq4, 2lr0, 2lrh, 2lse, 2lta, 2lv8, 2lvb, 2mbl, 2mbm, 2mn4, 2mq8, 2mra, 2mtt, 2mtu, 2muz, 2n1e, 2n2t, 2n2u, 2n3z, 2n41, 2n4e, 2n4n, 2n6h, 2n6i, 2n75, 2n76, 2n7n, 2n7o, 2n7t, 2n8d, 2nbl, 2nd2, 2nd3, 2o6n, 2rt4, 2rvd, 2x6p, 2zta, 3al1, 3cay, 3h5f, 3h5g, 3he4, 3he5, 3j89, 3kd7, 3ljm, 3ni3, 3ovj, 3ow9, 3p46, 3pbj, 3r3k, 3r46, 3r47, 3r48, 3r4a, 3r4h, 3ra3, 3s0r, 3t4f, 3tq2, 3twe, 3twf, 3twg, 3u29, 3v86, 3vjf, 3wn8, 4dac, 4dzk, 4dzl, 4dzm, 4dzn, 4e0k, 4e0l, 4e0m, 4e0n, 4e0o, 4g1a, 4g3b, 4g4l, 4g4m, 4h7r, 4h8f, 4h8g, 4h8l, 4h8m, 4h8o, 4hb1, 4ivh, 4ntp, 4ntr, 4nw8, 4nw9, 4oyd, 4p4v, 4p4w, 4p4x, 4p4y, 4p4z, 4p6j, 4p6k, 4p6l, 4pn8, 4pn9, 4pna, 4pnb, 4pnd, 4qtr, 4r80, 4tql, 4tut, 4uby, 4ubz, 4uos, 4uot, 4w5l, 4w5m, 4w5p, 4w5y, 4w67, 4w71, 4wbu, 4wbv, 4yxx, 4yxz, 4yy2, 4yy5, 4z1r, 5awl, 5bvl, 5byo, 5cwb, 5cwc, 5cwd, 5cwf, 5cwg, 5cwh, 5cwi, 5cwj, 5cwk, 5cwl, 5cwm, 5cwn, 5cwo, 5cwp, 5cwq, 5ehb, 5eoj, 5eon, 5et3, 5ez8, 5ez9, 5eza, 5ezc, 5eze, 5f2y, 5hkn, 5hkr, 5izs, 5j0h, 5j0i, 5j0j, 5j0k, 5j0l, 5j10, 5j2l, 5j73, 5jg9, 5jhi, 5ji4, 5jqz, 5k7v, 5k92, 5kb0, 5kb1, 5kb2, 5kpe, 5kph, 5kvn, 5kwo, 5kwp, 5kwx, 5kwz, 5kx0, 5kx1, 5kx2, 5l33, 5lo4, 5sbg, 5sbi, 5sbj, 5tgw, 5tgy, 5tph, 5tpj, 5trv, 5ts4, 5tx8, 5u35, 5u59, 5u5a, 5u5b, 5u5c, 5u9t, 5u9u, 5ugk, 5uoi, 5up5, 5uxt, 5v2g, 5v2o, 5v63, 5v64, 5v65, 5vli, 5vmr, 5vsg, 5vte, 5w0j, 5w9f, 5wlj, 5wlk, 5wll, 5wlm, 5woc, 5wod, 5wrx, 5yan, 6anf, 6anm, 6ann, 6b17, 6b87, 6c2u, 6c2v, 6c4x, 6c4y, 6c4z, 6c50, 6c51, 6c52, 6cfa, 6czg, 6czh, 6czi, 6czj, 6d02, 6d0t, 6dkm, 6dlc, 6dlm, 6dm9, 6dma, 6dmp, 6e5c, 6e5h, 6e5i, 6e5j, 6e5k, 6egc, 6egl, 6egm, 6egn, 6ego, 6egp, 6eik, 6eiz, 6fce, 6g65, 6g66, 6g67, 6g68, 6g69, 6g6a, 6g6b, 6g6c, 6g6d, 6g6e, 6g6f, 6g6g, 6g6h, 6hqe, 6i1j, 6jcc, 6kos, 6m6z, 6mcd, 6mct, 6mpw, 6mq2, 6mqu, 6mrr, 6mrs, 6msq, 6msr, 6n4n, 6n9h, 6naf, 6nx2, 6nxm, 6ny8, 6nye, 6nyi, 6nyk, 6nz1, 6nz3, 6o0c, 6o0i, 6o35, 6o3n, 6ohh, 6oln, 6olo, 6os8, 6osd, 6ov9, 6ovs, 6ovu, 6ovv, 6owd, 6q1w, 6q22, 6q25, 6q5h, 6q5i, 6q5j, 6q5k, 6q5l, 6q5m, 6q5n, 6q5o, 6q5p, 6q5q, 6q5r, 6q5s, 6r28, 6s3d, 6tms, 6tt6, 6tvj, 6u1s, 6u47, 6ucx, 6ud9, 6udr, 6udw, 6udz, 6uf4, 6uf7, 6uf8, 6uf9, 6ufa, 6ufu, 6ug2, 6ug3, 6ug6, 6ugb, 6ugc, 6v4y, 6v50, 6v57, 6v58, 6v5g, 6v5i, 6v5j, 6v67, 6v8e, 6vzx, 6w3w, 6w40, 6w46, 6w47, 6w6x, 6w70, 6wi5, 6wmk, 6wrv, 6wvs, 6x8n, 6xeh, 6xwi, 6xxz, 6xy0, 6xy1, 6yaz, 6yb0, 6yb1, 6yb2, 6yqx, 6yqy, 6z0l, 6z0m, 6zt1, 7a1t, 7a8s, 7ah0, 7arr, 7ars, 7bas, 7bat, 7bau, 7bav, 7baw, 7bey, 7bim, 7bo8, 7bo9, 7boa, 7bpl, 7bpm, 7bpn, 7bpp, 7bqb, 7bqc, 7bqd, 7bqe, 7bqm, 7bqn, 7bqq, 7bqs, 7c0n, 7cbc, 7dkk, 7dko, 7dns, 7fao, 7jh5, 7kuw, 7l33, 7ldf, 7lib, 7lmv, 7lmx, 7lxp, 7lxq, 7m5t, 7mcc, 7mcd, 7nff, 7nfg, 7nfh, 7nfi, 7nfj, 7nfk, 7nfl, 7nfm, 7nfn, 7nfo, 7nfp, 7osu, 7ot7, 7q1q, 7q1r, 7q1s, 7q1t, 7qdi, 7qdj, 7qdk, 7qwa, 7qwb, 7qwc, 7qwd, 7qwe, 7rkc, 7rmx, 7skn, 7sko, 7skp, 7smj, 7sq3, 7sq4, 7sq5, 7t6e, 7tjl, 7tls, 7tlu, 7tm1, 7tm2, 7tma, 7tme, 7tmh, 7tmi, 7tmj, 7tmk, 7tml, 7ubc, 7ubd, 7ube, 7ubf, 7ubg, 7ubh, 7ubi, 7ucp, 7udk, 7udl, 7udm, 7udn, 7udo, 7udv, 7udw, 7udx, 7udy, 7udz, 7uek, 7uit, 7unh, 7uni, 7upo, 7upp, 7upq, 7ur7, 7ur8, 7uuq, 7uwy, 7uwz, 7uzl, 7y9c, 7yh8, 7zbs, 8a09, 8a3g, 8a3i, 8a3j, 8a3k, 8a50, 8a51, 8ah9, 8ang, 8anh, 8ani, 8ank, 8anm, 8b15, 8b16, 8b45, 8bcs, 8bct, 8bfd, 8bfe, 8c3e, 8c3w, 8cto, 8cun, 8cwa, 8cyk, 8d03, 8d04, 8d05, 8d06, 8d07, 8d08, 8d09, 8ddf, 8ddg, 8ddh, 8dpy, 8dt0, 8e55, 8ec9, 8eca, 8ek4, 8eov, 8eox, 8eoz, 8erw, 8evm, 8f4x, 8f53, 8f54, 8fbn, 8fg6, 8fih, 8fje, 8fjf, 8fjg, 8fvt, 8g8i, 8ga6, 8ga7, 8ga9, 8gaa, 8gaq, 8gb9, 8gba, 8gbh, 8gbi, 8gbm, 8gbo, 8gd6, 8gd8, 8giv, 8gj7, 8gjc, 8gjd, 8gk1, 8gk2, 8gk9, 8gkb, 8gkx, 8gl0, 8gl3, 8gl4, 8gl5, 8h7c, 8h7d, 8h7e, 8hvs, 8i8y, 8ju8, 8k7m, 8k7o, 8k7z, 8k83, 8k84, 8k8f, 8k8g, 8k8i, 8ka6, 8ka7, 8kac, 8kc0, 8kc1, 8kc4, 8kc5, 8kc8, 8kcj, 8kck, 8kdq, 8oh2, 8ohi, 8ohp, 8oi0, 8onq, 8oys, 8oyv, 8oyw, 8oyx, 8oyy, 8qaa, 8qab, 8qac, 8qag, 8qah, 8qai, 8qkd, 8sw2, 8sy4, 8szz, 8t5e, 8t61, 8t62, 8t63, 8tn1, 8tn6, 8tnb, 8tnc, 8tnd, 8tnm, 8tno, 8txs, 8un8, 8utx, 8v56, 8v59, 8v5w, 8v5x, 8v5z, 8v61, 8vc8, 8vei, 8vej, 8vl3, 8vl4, 8vog, 8vp7, 8vpc, 8vpd, 8vpe, 8vpj, 8vps, 8vpt, 8vpx, 8vpy, 8vpz, 8vq0, 8vsf, 8vt8, 8vw7, 8vw8, 8vx |
| MMseqs2 output and analysis results | 1abz, 1coi, 1cos, 1ec5, 1jm0, 1jmb, 1le1, 1lq7, 1nvo, 1qys, 1tjb, 1uw1, 2a3d, 2gjh, 2jgo, 2jvf, 2kl8, 2kpo, 2l69, 2l82, 2lnd, 2lq4, 2lv8, 2lvb, 2mbl, 2mbm, 2mra, 2n3z, 2n41, 2n4e, 2n75, 2x6p, 2zta, 3h5f, 3h5g, 3he5, 3j89, 3ljm, 3pbj, 4hb1, 4oyd, 4tql, 4yxx, 4yxz, 5bvl, 5byo, 5cwb, 5cwc, 5cwd, 5cwg, 5cwh, 5cwi, 5cwj, 5cwk, 5cwl, 5cwm, 5cwn, 5cwo, 5cwp, 5cwq, 5j0h, 5j0i, 5jg9, 5k92, 5kb0, 5kb1, 5kb2, 5lo4, 5tgw, 5tgy, 5tph, 5tpj, 5trv, 5u35, 5u9t, 5u9u, 6c2u, 6c2v, 6egl, 6egm, 6egn, 6ego, 6egp, 6mcd, 6mct, 6mpw, 6mq2, 6msq, 6msr, 6n4n, 6n9h, 6naf, 6nye, 6os8, 6osd, 6ov9, 6ovu, 6s3d, 6v5g, 6v5i, 6v8e, 6w6x, 6w70, 6wrv, 6wvs, 6x8n, 6xwi, 6yqx, 6yqy, 7a8s, 7ah0, 7fao, 7lmv, 7lmx, 7mcc, 7mcd, 7osu, 7ot7, 7skn, 7sko, 7skp, 7smj, 7udv, 7udw, 7udx, 7udy, 7udz, 7yh8, 8a50, 8a51, 8c3e, 8c3w, 8evm, 8t5e, 8tn1, 8tn6, 8tnb, 8tnc, 8tnd |
| Foldseek input and analysis results | 1abz, 1al1, 1bb1, 1byz, 1coi, 1cos, 1d7t, 1djf, 1fme, 1fsd, 1fsv, 1g6u, 1hcw, 1hqj, 1j4m, 1jy6, 1jy9, 1k43, 1kyc, 1l4x, 1le0, 1le1, 1lq7, 1m3w, 1n09, 1n0a, 1n0c, 1n0d, 1pbz, 1pyz, 1qp6, 1rh4, 1s9z, 1sn9, 1sna, 1sne, 1t8j, 1tgg, 1tjb, 1u0i, 1uno, 1vl3, 1vm5, 1vrz, 1xof, 2a3d, 2evq, 2gjh, 2jgo, 2jvf, 2k6r, 2kjn, 2kjo, 2klw, 2l96, 2l99, 2l9a, 2lny, 2lq4, 2mtt, 2mtu, 2muz, 2n4n, 2n63, 2n6h, 2n6i, 2n7n, 2n7o, 2n7t, 2n8d, 2nbl, 2o6n, 2rt4, 2rvd, 2x6p, 2zta, 3al1, 3cay, 3h5f, 3h5g, 3he4, 3he5, 3j89, 3ljm, 3ni3, 3ovj, 3ow9, 3p46, 3pbj, 3r3k, 3r46, 3r47, 3r48, 3r4a, 3r4h, 3ra3, 3s0r, 3t4f, 3tq2, 3twe, 3twf, 3twg, 3u29, 3v86, 3vjf, 3wn8, 4dac, 4dzk, 4dzl, 4dzm, 4dzn, 4e0k, 4e0l, 4e0m, 4e0n, 4e0o, 4g1a, 4g3b, 4g4l, 4g4m, 4hb1, 4ivh, 4ntp, 4ntr, 4nw8, 4nw9, 4p6j, 4p6k, 4p6l, 4pn8, 4pn9, 4pna, 4pnb, 4pnd, 4tut, 4uby, 4ubz, 4uot, 4w5l, 4w5m, 4w5p, 4w5y, 4w67, 4w71, 4wbu, 4wbv, 4z1r, 5awl, 5ehb, 5et3, 5ez8, 5ez9, 5eza, 5ezc, 5eze, 5f2y, 5hkn, 5hkr, 5k92, 5kb0, 5kb1, 5kb2, 5kwp, 5kwx, 5kx0, 5kx1, 5kx2, 5mfe, 5u59, 5u5a, 5u5b, 5u5c, 5u9t, 5u9u, 5ugk, 5uxt, 5v2g, 5v63, 5v64, 5vsg, 5w0j, 5wlj, 5wlk, 5wll, 5wlm, 5woc, 5wrx, 5yan, 6anf, 6anm, 6ann, 6b17, 6c4x, 6c4y, 6c4z, 6c50, 6c51, 6c52, 6cfa, 6d02, 6dm9, 6e5i, 6e5j, 6e5k, 6egl, 6egm, 6egn, 6ego, 6egp, 6eik, 6eiz, 6fce, 6g65, 6g66, 6g67, 6g68, 6g69, 6g6a, 6g6b, 6g6c, 6g6d, 6g6e, 6g6f, 6g6g, 6g6h, 6mcd, 6mct, 6mpw, 6mq2, 6mqu, 6o3n, 6oln, 6olo, 6os8, 6ov9, 6ovs, 6ovu, 6ovv, 6owd, 6q1w, 6q22, 6q25, 6r28, 6tt6, 6tvj, 6u47, 6ucx, 6ud9, 6udr, 6udw, 6udz, 6uf4, 6uf7, 6uf8, 6uf9, 6ufa, 6ufu, 6ug2, 6ug3, 6ug6, 6ugb, 6ugc, 6v4y, 6v50, 6v57, 6v58, 6v5g, 6v5j, 6vzx, 6w46, 6w47, 6wky, 6wl0, 6wl1, 6xxz, 6xy1, 6yaz, 6yb0, 6yb1, 6yb2, 6zt1, 7a1t, 7bas, 7bau, 7bav, 7baw, 7bo8, 7bo9, 7boa, 7c0n, 7eq9, 7l33, 7lxp, 7lxq, 7nff, 7nfg, 7nfh, 7nfi, 7nfj, 7nfk, 7nfl, 7nfm, 7nfn, 7nfo, 7nfp, 7q1q, 7q1r, 7q1s, 7q1t, 7qdi, 7qdj, 7qdk, 7qnl, 7qnp, 7qwa, 7qwb, 7qwc, 7qwd, 7qwe, 7rx5, 7t6e, 7tls, 7tlu, 7tm1, 7tm2, 7tma, 7tme, 7tmh, 7tmi, 7tmj, 7tmk, 7tml, 7ubc, 7ubd, 7ube, 7ubf, 7ubg, 7ubh, 7ubi, 7ucp, 7udm, 7udv, 7udw, 7udx, 7udy, 7udz, 7uit, 7uuq, 7uzl, 7xm1, 7zbs, 8a09, 8a3i, 8a3k, 8a50, 8ang, 8anh, 8ani, 8ank, 8anm, 8b15, 8b16, 8b45, 8bfd, 8bfe, 8cto, 8cun, 8cwa, 8ddf, 8ddg, 8ddh, 8dpy, 8ec9, 8eca, 8f53, 8gb9, 8gba, 8gbh, 8gbi, 8gbm, 8gbo, 8gd6, 8gd8, 8giv, 8gj7, 8gjd, 8gk1, 8gk2, 8gk9, 8gkb, 8gkx, 8gl0, 8gl4, 8gl5, 8glt, 8hvs, 8oh2, 8ohi, 8ohp, 8oi0, 8onq, 8qye, 8sw2, 8sy4, 8t61, 8t62, 8t63, 8umr, 8un1, 8un8, 8utx, 8v56, 8v59, 8v5w, 8v5x, 8v5z, 8v61, 8vog, 8vp7, 8vpc, 8vpd, 8vpe, 8vpj, 8vps, 8vpt, 8vpx, 8vpy, 8vpz, 8vq0, 8vsf, 8vt8, 8vw7, 8vw8, 8vxs |
| Foldseek output and analysis results | 1abz, 1al1, 1bb1, 1coi, 1cos, 1djf, 1fme, 1fsd, 1fsv, 1g6u, 1hcw, 1j4m, 1jy6, 1jy9, 1k43, 1le0, 1le1, 1lq7, 1m3w, 1n0a, 1n0c, 1n0d, 1pyz, 1qp6, 1rh4, 1s9z, 1sn9, 1sna, 1sne, 1t8j, 1tgg, 1tjb, 1u0i, 1vm5, 1vrz, 1xof, 2a3d, 2evq, 2jgo, 2k6r, 2kjn, 2kjo, 2klw, 2l96, 2l99, 2l9a, 2lny, 2lq4, 2mtt, 2mtu, 2muz, 2n4n, 2n63, 2n6h, 2n6i, 2n8d, 2nbl, 2o6n, 2rt4, 2rvd, 2x6p, 2zta, 3h5f, 3h5g, 3he4, 3he5, 3j89, 3ljm, 3pbj, 3r3k, 3r46, 3r47, 3r48, 3r4a, 3r4h, 3ra3, 3s0r, 3tq2, 3twe, 3twf, 3twg, 3v86, 3vjf, 4dac, 4dzk, 4dzl, 4dzm, 4dzn, 4e0l, 4e0m, 4e0n, 4e0o, 4g1a, 4g3b, 4g4l, 4g4m, 4hb1, 4ivh, 4ntp, 4ntr, 4nw8, 4nw9, 4p6j, 4p6k, 4p6l, 4pn8, 4pn9, 4pna, 4pnb, 4pnd, 4uot, 5awl, 5ehb, 5et3, 5ez8, 5ez9, 5eza, 5ezc, 5eze, 5f2y, 5hkn, 5hkr, 5k92, 5kb0, 5kb1, 5kb2, 5kwp, 5kwx, 5kx0, 5kx1, 5kx2, 5u59, 5u5a, 5u5b, 5u5c, 5u9t, 5u9u, 5uxt, 5v2g, 5v63, 5v64, 5w0j, 5wlj, 5wlk, 5wll, 5wlm, 5woc, 5wrx, 5yan, 6anf, 6b17, 6c4x, 6c4y, 6c4z, 6c51, 6c52, 6cfa, 6d02, 6dm9, 6e5i, 6e5j, 6e5k, 6egl, 6egm, 6egn, 6ego, 6egp, 6eik, 6eiz, 6g68, 6g69, 6g6a, 6g6b, 6g6c, 6g6d, 6g6e, 6g6f, 6g6g, 6g6h, 6mcd, 6mct, 6mpw, 6mq2, 6mqu, 6o3n, 6oln, 6olo, 6os8, 6ov9, 6ovs, 6ovu, 6ovv, 6owd, 6q1w, 6q22, 6q25, 6r28, 6tvj, 6u47, 6udw, 6ufa, 6ug6, 6v4y, 6v50, 6v57, 6v58, 6v5g, 6v5j, 6wky, 6wl0, 6wl1, 6xxz, 6xy1, 6yaz, 6yb0, 6yb1, 6yb2, 6zt1, 7a1t, 7bas, 7bau, 7bav, 7baw, 7bo8, 7bo9, 7boa, 7l33, 7nff, 7nfg, 7nfh, 7nfi, 7nfj, 7nfk, 7nfl, 7nfm, 7nfn, 7nfo, 7nfp, 7q1q, 7q1r, 7q1s, 7q1t, 7qdj, 7qdk, 7qwa, 7qwb, 7qwc, 7qwd, 7qwe, 7rx5, 7udv, 7udw, 7udx, 7udy, 7udz, 7zbs, 8a09, 8a3i, 8a50, 8b15, 8b16, 8b45, 8bfe, 8dpy, 8hvs, 8t61, 8t62, 8t63 |

Table S4: PDB codes missing between different steps of similarity analysis. These designs are likely to show very high degree of novelty.

#### Software versions:

MMseqs2 version: ad5837b3444728411e6c90f8c6ba9370f665c443

Foldseek version: 9.427df8a

DE-STRESS version: 3d6c2419bc1f7481fd0c4a5d933fcdb535aca457

#### Frontend requirements:

<https://elm-lang.org/> and <https://elm.land/>

"NoRedInk/elm-json-decode-pipeline": "1.0.1",

"elm/browser": "1.0.2",

"elm/core": "1.0.5",

"elm/html": "1.0.0",

"elm/http": "2.0.0",

"elm/json": "1.1.3",

"elm/svg": "1.0.1",

"elm/time": "1.0.0",

"elm/url": "1.0.0",

"elm-community/list-extra": "8.7.0",

"feathericons/elm-feather": "1.5.0",

"gicentre/elm-vega": "5.7.1",

"justinmimbs/date": "4.1.0",

"krisajenkins/remotedata": "6.0.1",

"mdgriffith/elm-ui": "1.1.8",

"miniBill/elm-codec": "2.1.0"

#### Backend requirements:

"click==8.1.7",

"Flask==3.0.3",

"Flask-Cors==3.0.9",

"pymongo==4.7.2",

<https://github.com/tiangolo/meinheld-gunicorn-docker>

<https://hub.docker.com/_/mongo>
